# Single cell multi-omics reveals early elevated function and multiple fates within human progenitor exhausted CD8^+^ T cells

**DOI:** 10.1101/2021.09.09.459584

**Authors:** Curtis Cai, Jerome Samir, Mehdi R. Pirozyan, Thiruni N. Adikari, Money Gupta, Preston Leung, Brendan Hughes, Willem Van der Byl, Simone Rizzetto, Auda Elthala, Elizabeth Keoshkerian, Jean-Louis Palgen, Timothy Peters, Thi H. O. Nguyen, Raymond Louie, Katherine Kedzierska, Silvana Gaudieri, Rowena A. Bull, Andrew R. Lloyd, Fabio Luciani

**Author notes:** Corresponding author: Associate Professor Fabio Luciani Immunogenomics laboratory, School of Medical Sciences, and the Kirby Institute University of New South Wales Sydney 2052 Australia. Equally contributed (Curtis Cai, Jerome Samir).

## Abstract

T-cell exhaustion is a hallmark of hepatitis C virus (HCV) infection and limits protective immunity in chronic viral infections and cancer. Limited knowledge exists of the initial viral and immune dynamics that characterise exhaustion in humans. We studied longitudinal blood samples from a unique cohort of subjects with primary infection using single cell multi-omics to identify the functions and phenotypes of HCV-specific CD8^+^ T cells. Early elevated IFN-γ response against the transmitted virus was associated with the rate of immune escape, larger clonal expansion, and early onset of exhaustion. Irrespective of disease outcome we discovered progenitors of early-exhaustion with intermediate expression of PD-1. Intra clonal analysis revealed distinct trajectories with multiple fates suggesting evolutionary plasticity of precursor cells. These findings challenge current paradigm on the contribution of CD8^+^ T cells to HCV disease outcome and provide data for future studies on T-cell differentiation in human infections.

**One sentence summary:** Progenitors of T-cell exhaustion in acute HCV infection

## INTRODUCTION

The cytotoxic CD8^+^ T-cell response during primary viral infection involves recruitment, expansion, and differentiation of epitope-specific clones from the naïve population. The selective pressure exerted by these responses on rapidly mutating RNA viruses, may drive the generation of immune escape mutations(*1*). In chronic infections, epitope-specific T-cell responses may acquire an exhausted phenotype after sustained antigen exposure, which is characterised by reduced responsiveness to further stimulation(*2*). Pioneering work in the murine lymphocytic choriomeningitis virus (LCMV) model, has mapped the molecular and phenotypic profiles of CD8^+^ T cells with acute resolving and chronic infections(*3–5*), revealing progenitors of exhaustion (T_PEX_), defined by the expression of transcription factors, TOX and TCF1, which arise in the acute phase of infection and sustain terminally exhausted subsets over the long-term(*6, 7*). A similar subset of progenitor has been identified within human memory populations (*8*), although how these cells arise and from which antigen specificity remains poorly defined.

Study on human T cell responses to chronic viral infections are limited by the scarcity of samples from the acute phase as many infections progress asymptomatically. Furthermore, the diversity of epitope targets, host HLA heterogeneity, and viral evolution require comprehensive analysis of immunological and virological variables to successfully identify individual-specific responses and their molecular and phenotypic profiles. Hepatitis C virus (HCV) infection is an excellent immunological model for understanding the development of exhaustion because both spontaneous clearance and chronic infections are naturally observed. The current model of the role of cytotoxic CD8^+^ T cells in contributing to HCV disease outcome assumes that along with host factors, T cell’s contributions to disease resolution are mediated by strong responses to a broad range of epitopes(*9, 10*), (*11*). Recent studies on antigen removal via antiviral therapy against HCV or following immune escape described a scar on the recovered memory CD8^+^ T-cell responses, suggesting that exhaustion may not be fully reversible (*6, 7*).

While human studies on the late phase of HCV infection greatly advanced our understanding on T cell exhaustion, how the early viral dynamics and magnitude of responses determine immune escape and early differentiation from progenitor cells remains incompletely characterised. In this study, we sampled longitudinal blood samples from a unique cohort of patients with primary HCV infections (*12, 13*) within weeks of transmission and through their peak of viremia until determination of clearance or chronic infection. We applied deep sequencing of viral populations, functional, and single cell multiomics to characterise viral dynamics and HCV-specific CD8^+^ T-cell responses. We found that the magnitude of IFN-γ responses during acute phase of chronic infection was associated with early differentiation into exhaustion and positively correlated with clonal expansion and rate of immune escape. We identified a progenitor state of exhausted cells, irrespective of disease outcome, which persisted after viral clearance. Our data contribute to understanding when and how exhausted T cell populations are formed and suggest a revision of the current paradigm of the role of CD8^+^ T cells in determining outcome of infection.

## RESULTS

### High magnitude of IFN-γ response is associated with rapid viral immune escape

We applied a comprehensive combination of assays to study viral evolution and HCV-specific CD8^+^ T-cell responses from longitudinal samples during early acute and chronic HCV infection (Fig. 1A-B). Circulating viral genomes were deep sequenced from both chronic progressors (CH) and clearers (CL) (table S1) to predict Human Leukocyte Antigen class I (HLA-I) epitopes and validated by IFN-γ ELISpot (Fig. 1C, table S2). We identified 50 epitopes across 14 subjects (table S3). Notably, 20 epitopes acquired amino acid mutations, and all were detected in chronic progressors (Fig. 1D). Interestingly, the breadth (i.e., number of unique epitope specific responses) and magnitude of IFN-γ responses from chronic progressors were higher than clearers, up to the first 120 days post-infection (DPI) (Fig. 1E). During this early phase, there were no significant differences in levels of viral load (first 120 DPI, Fig. 1A), which is the timeframe when acute viremia is successfully controlled by spontaneous clearers and rebound in chronic progressors(*14*). Notably, the magnitude of responses rapidly declined over time in chronic progressors, while these were sustained in clearers.

**Fig. 1.**
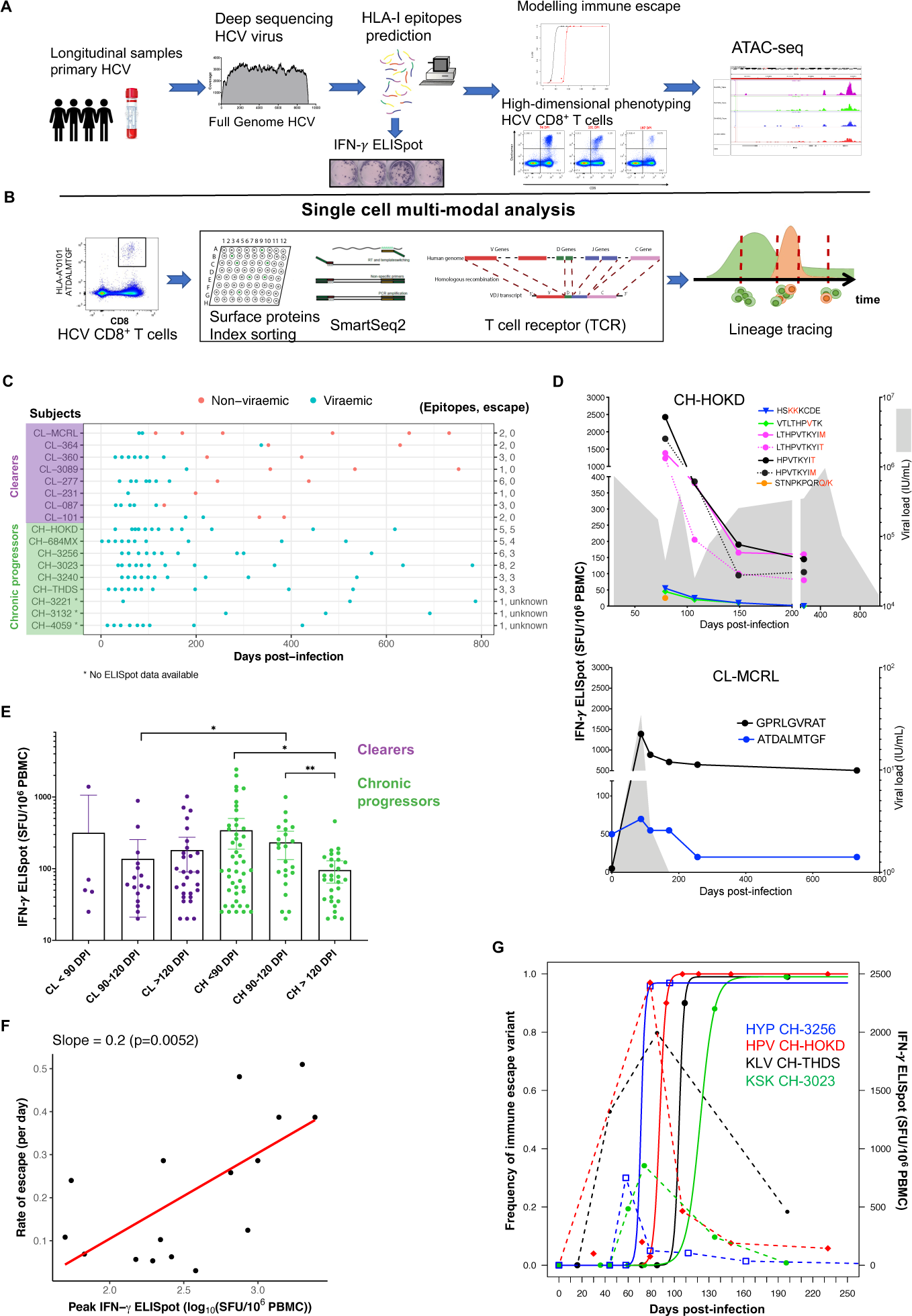
Comprehensive analyses reveal early and functional HCV-specific CD8^+^ T cells associated with viral immune escape. **(A, B)** Schematic of experimental and bioinformatic analyses used in this study. Viral genomes were sequenced from longitudinal blood samples from individuals with primary HCV infection and mutated variants were detected. HLA-I epitopes were predicted from sequence data and epitope-specific CD8^+^ T cells identified by IFN-γ ELISpot and dextramer staining. ATAC-seq was performed on HCV-specific and other CD8+ T cells. HCV-specific populations were analysed by flow cytometry, single-cell RNA-sequencing and for paired T-cell receptor sequences. **(C)** Swimming plot outlining subjects and sampling timepoints. Colours represent viraemic and non-viraemic sample timepoints. Legend shows the total number of epitopes, and those that escape for each subject. CL: clearers, CH: chronic progressors. **(D)** Kinetics of viral load (shaded areas, measured in international units (IU)) and IFN-production (Spot forming unit (SFU/Million PBMC)) for epitope-specific CD8^+^ T-cell responses in two representative subjects plotted over days post-infection. Lines represent distinct epitope-specific responses: solid lines indicate responses targeting wildtype epitope (from the transmitted founder sequence) and dashed lines indicate those targeting mutated epitope (identified from amino acid mutations within the wild-type epitope over the course of the infection, see Supplementary note 1); amino acid mutations are coloured in red. **(E)** Comparison of IFN-γ responses according to the sample timepoint and disease outcome. Each point represents a unique subject, epitope, and timepoint combination. Ranges show median and 95% confidence interval. Comparisons were performed using Mann-Whitney test. * p < 0.05, ** p < 0.01. CL: clearers, CH: chronic progressors. **(F)** Scatter plot revealing a significant correlation between the estimated rates of immune escape of target epitopes and the maximum IFN-γ ELISpot responses (log_10_ transformed) measured against the corresponding epitope within the first 120 DPI. Only subjects who progressed to chronic infection and with available longitudinal deep sequencing data of viral populations were included. Data were fitted with a linear regression and coefficient of slope is reported. **(G)** Kinetics of viral immune escape variants using estimated rates (continuous lines) from the frequency of vial escape variants measured from deep sequencing of viral genome, and IFN-γ ELISpot levels (Dashed lines connect the measured SFU values) for four epitope-specific populations.

We next quantified the rate at which viral epitope undergo immune escape by fitting the frequency of amino acid mutations within epitopes to a model of virus evolution with T cell killing (see Supplementary Methods). We observed a positive correlation between the rate of escape and the peak-value of IFN-γ production against the same epitopes (Fig. 1F). Estimated rates ranged from 0.03 to 0.51 day^-1^ (table S5), which implied a broad time scale of 28-460 days for the observed escape variants to comprise over half of the total viral population. Notably, the earliest peak of IFN-γ responses measured in this study preceded the estimated time at which escape variants became dominant (Fig. 1G), thus suggesting that early onset of high magnitude response contribute to rapid immune escape. Notably, in three subjects a reduced IFN-γ response was measured against the mutated epitope sequences (CH-3023, CH-684MX, CH-3240), thus suggesting persisting T cell response against immune escape (Fig. 1D, table S10). These results show that the IFN-γ producing cells occur earlier and in higher number in chronic progressors, but rapidly decline over time irrespective of the occurrence of immune escape.

### Subsets of progenitors and exhaustion during acute phase infection

We next investigated the phenotype of HCV-specific responses over the course of the infection. We utilised dextramers to study longitudinally the phenotype of 20 epitope-specific CD8^+^ T cell responses that were identified in the previous experiment in five clearers and seven chronic progressors (table S4). There was a strong correlation between the size of HCV-specific populations detected by IFN-γ ELISpot and dextramer staining. Notably, larger populations were detected by dextramer staining which suggested that not all cells were functionally responsive (Fig. S1B). T-cell phenotypes were assessed by co-expression of CD127 and PD-1 to distinguish between exhausted PD-1^high^CD127^low^ (TEX), memory-like PD-1^high^CD127^high^ (T_ML_), memory PD-1^low^ CD127^high^ (T_MEM_), and effector PD-1^low^CD127^low^ subsets (Fig. 2A-B). We observed a subtype of PD-1^int^CD127^low^, herein referred to as progenitor of exhaustion (T_PEX_, see transcriptional signature analysis below), which was larger than the T_EX_ subset in both chronic progressors and clearers subjects (Fig. 2C). Additional surface and intracellular markers revealed in chronic progressors increased proportions of T-bet^+^, T-bet^+^Eomes^-^, CTLA4^+^, KLRG1^+^, Tim-3^+^ and CD38^+^ phenotypes (Fig. 2D, Fig. S1), and lower proportions of T_MEM_ and T_ML_ subsets. Clearers and chronic progressors did not significantly differ in the proportion of T_EX_ and T_PEX_, nor in CD160^+^ and 2B4^+^ subsets (Fig. 2D, Fig. S1).

**Fig. 2.**
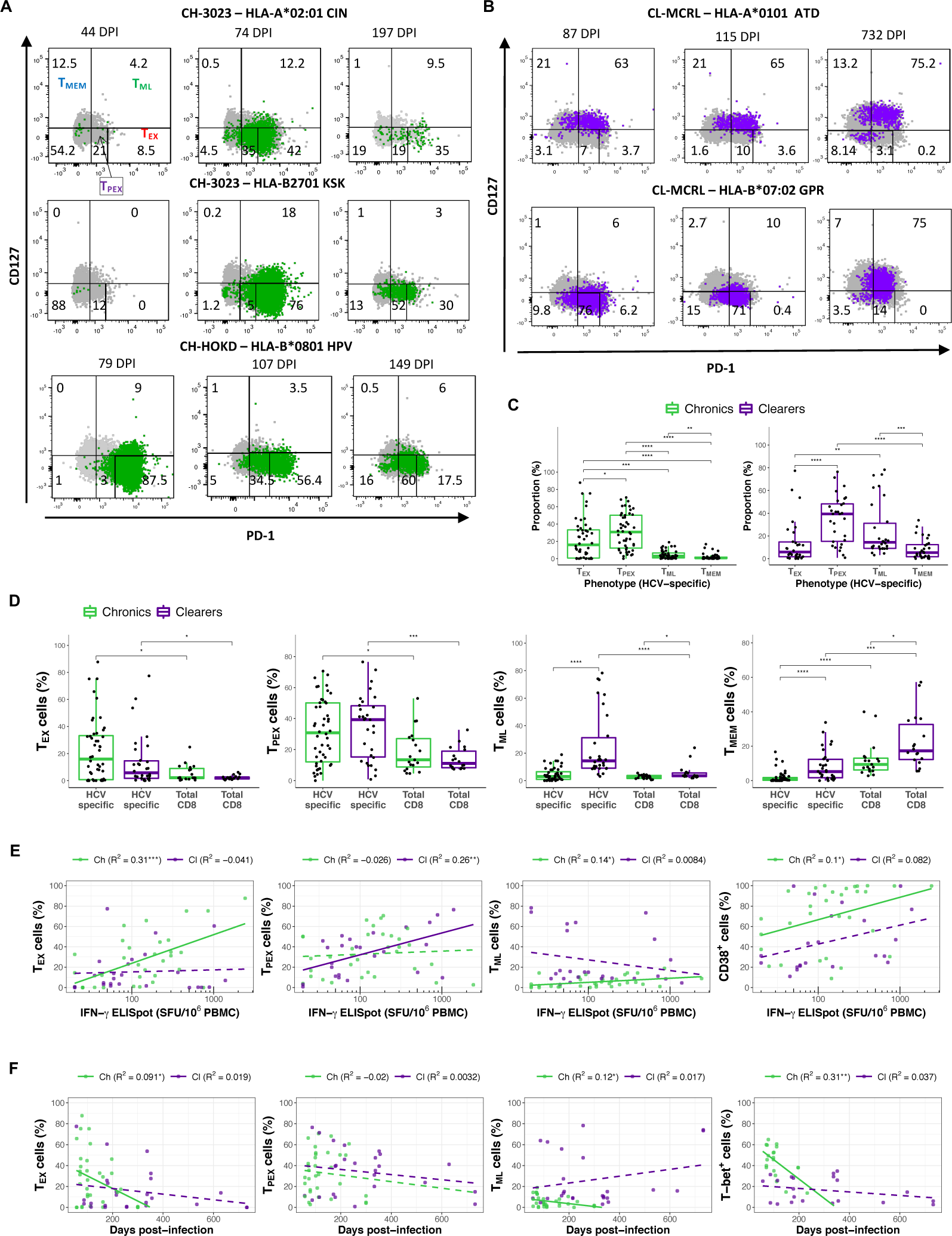
Subsets of progenitors and exhaustion of HCV-specific CD8^+^ T cells during acute phase infection. **(A-B)** Representative flow cytometry gating plots of dextramer-positive CD8^+^ T cell populations analysed longitudinally targeting autologous epitopes identified from circulating viral populations, in chronic progressors (A) and clearers (B). Each plot shows the frequency of dextramer positive cells based on the expression of CD127 and PD-1 expression across five distinct subsets: T_EX_/exhausted (PD-1^high^CD127^low^), T_PEX_/progenitor-exhausted (PD-1^int^CD127^low^), T_ML_/memory-like (PD-1^high^CD127^high^), T_MEM_/memory (PD-1^low^CD127^high^), and effector (CD127^low^PD-1^low^). DPI: days post-infection, HLA: human leukocyte antigen. Shown are cells gated from a lymphocyte/singlet/live population that were CD19 negative, CD3, and CD8 positive. **(C)** Comparison of phenotypic subsets of HCV-specific populations defined by PD-1 and CD127 co-expression, segregated by disease outcome. Individual points represent unique subject, timepoint and epitope specificity. **(D)** Comparison (box plots) of the proportion of cells from each population with positive expression for markers measured by flow cytometry compared between HCV-specific CD8^+^ T cells in both disease outcomes, and total CD8^+^ T cells. **(E)** Scatter plots showing the relation between the proportion of dextramer positive CD8^+^ T-cell populations with positive expression of phenotypic markers (measured by flow cytometry) and IFN-γ ELISpot values (number of SFU per million PBMC). Lines represent linear regression, adjusted coefficients and p-values are shown in the legend of each plot. Lines are dashed if insignificant (p-value > 0.05). **(F)** Same as for (E) but proportions are plotted and regressed against DPI in place of IFN-γ ELISpot. Statistical comparisons between groups were performed with Wilcoxon Rank Sum test (* p < 0.05, ** p < 0.01, *** p < 0.001, **** p < 0.0001).

We analysed the relationships between phenotypes of HCV-specific T cells in the context of their magnitude of IFN-γ production and time since infection by linear regressions. The number of IFN-γ producing T cells in chronic progressors increased with the proportion of T_EX_, T_ML_, and activated CD38^+^ cells (Fig. 2E, Fig. S1). In clearers, T_PEX_ increased with the IFN-γ producing T cells, as well as subsets expressing Eomes^+^, and exhaustion associated 2B4^+^, and CD160^+^. Over the course of the infection, the proportion of T_PEX_ remained stable in both disease outcomes (Fig. 2F, Fig. S1), while in chronic progressors a rapid decline of T_EX_, T-bet^+^ and CD38^+^ subsets were observed. Altogether, these results revealed that HCV-specific populations in chronic progressors are characterised by a larger expansion of IFN-γ producing cells, consistent with an early onset of exhausted-like cells, expressing T-bet, Tim-3, KLRG1, and CTLA-4, but no significant differences in PD-1 and exhausted-associated markers CD160, and 2B4 compared to clearers, which are characterised by an early and stable T_MEM_ phenotype.

### Single-cell multi-omics unveil molecular and phenotypic heterogeneity of HCV-specific CD8^+^ T cells

The identification of distinct subsets of PD-1 and CD127 expression within HCV-specific populations of CD8^+^ T cells demonstrated phenotypic heterogeneity which we explored further with single-cell RNA-sequencing (scRNA-seq) combined with clonal analysis using T-cell receptor (TCR) sequences, and protein expression from index sorting (Fig. 1B). We compiled a dataset of 1603 single cells passing quality control from nine subjects (seven chronic progressors and two clearers). Dimensionality reduction (UMAP) from scRNA-seq revealed a heterogeneous distribution of cells that was mostly explained by disease outcome, time point and magnitude of IFN-γ response of the sample of origin (Fig. 3A), and in part by subject or epitope specificity (fig. S2). Cells from chronic progressors had a higher expression of *TBX21*, *TOX*, *LCK* (T-cell signalling), *GZMK*, and *NKG7*, while those from clearers had a higher expression of memory-like associated genes (*IL7R*, *TCF7, FOS, JUN*) in both the early and late phases (Fig. 3B). We utilised protein and gene expression profiles to identify signatures of phenotypic subsets. T_PEX_ cells had an intermediate transcriptomic profile between exhaustion and memory subsets (Fig. 3C-D, table S8). When compared to T_EX_ cells, T_PEX_ had lower expression of exhaustion (*TOX*, *TIGIT*, *PDCD1*), cytotoxic genes (*NKG7*, *GZMH*, *PRF1*), and, like T_MEM_ and T_ML_, these cells expressed higher level of *IL7R*, *TCF7*, *FOS, JUNB,* and lower of *NFATC3*. These trends were also confirmed by protein index sorting (fig. S2C).

**Fig. 3.**
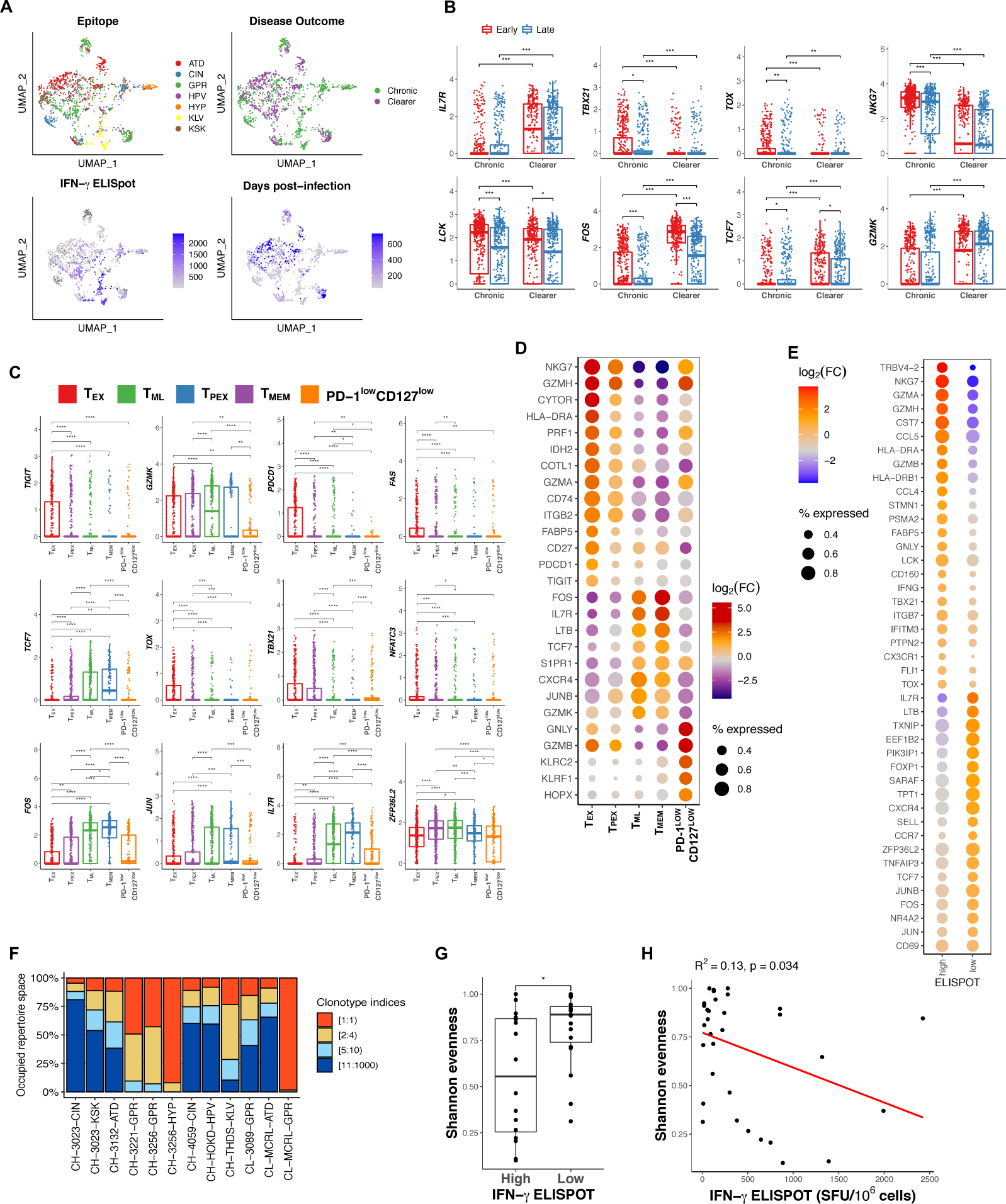
Single-cell multi-omics uncover molecular and phenotypic heterogeneity of HCV-specific CD8^+^ T cells. **(A)** Dimensionality reduction (UMAP) of scRNAseq data from HCV-specific CD8^+^ T cells (1603 cells). The four panels show cells coloured by epitope and infection characteristics as per legend. **(B)** Comparison (box plots) of gene expression levels (log_10_(TPM + 1)) for selected genes from individual cells grouped by infection phase (Early: before or equal to 120 DPI, late: after 120 DPI) and disease outcome. **(C)** Comparison (box plots) of gene expression levels for selected genes from individual cells grouped by phenotype based on co-expression of PD-1 and CD127. **(D)** Dot plot of selected genes (p < 0.05, |log_2_(FC)| ≥ 1) identified from differential gene expression analysis between cells grouped by phenotype. The ball size represents the proportion of cells with non-zero expression from each phenotype. The colour represents fold-change relative to all other cells from other phenotypes. **(E)** Dot plot of selected genes obtained from differentially expression analysis between cells grouped by magnitude of IFN-γ of the corresponding immune response (High: >211 SFU, 85^th^ percentile of the measured values; low IFN-γ ≤211 SFU). **(F)** Distribution (Stacked bar plot) of the size of clones identified from each epitope specific population in each subject. Colour legend indicates groups of clones ranked by their size, e.g. [2:4) indicates the proportion of sample occupied by the second, third and fourth largest clones. **(G)** Group comparison (box plot) of T Cell Receptor (TCR) diversity measured by Shannon evenness between high and low magnitude of IFN-γ responses. Individual points represent measurements from T cell populations defined by epitope specificity in each sample time point in each subject. (H) Scatter plot identifying a positive relationship between TCR diversity and magnitude of IFN-γ responses. A linear regression (Drawn in red) was applied to the scatter plot and R^2^ and p-value were reported. In (G-H) data includes single-cell TCR sequences from Sanger sequencing in addition to those obtained from scRNA-seq dataset (Detailed in Supplementary Note 1, and Supplementary table S7). Statistical comparisons between groups (B, C) were performed with Wilcoxon Rank Sum Test (* p < 0.05, ** p < 0.01, *** p < 0.001, **** p < 0.0001).

We reasoned that a variable gene expression profile would characterise cells associated with a significant difference in the associated magnitude of IFN-γ response. We stratified cells into high and low IFN-γ responses with a threshold of 211 (Spot forming units (SFU)/million PBMC - the 85^th^ percentile of recorded values. High IFN-γ producing cells expressed exhausted-like (*TOX, PTPN2, CD160)*, TCR signalling (*LCK*), and functional (*GZMA*, *GZMB*, *NKG7*, *IFNG*) genes (Fig. 3E), while low IFN-γ responses expressed memory-like profiles (*IL7R*, *CD69*, *TCF7*, *NR4A2*) and genes such as *TNFAIP3* encoding A20, associated with limiting inflammatory responses. Comparison with early and late phase of infection confirmed a rapid decline of exhausted- and cytotoxic-associated profiles in high IFN-γ responses (fig. S2D).

By reconstructing full-length TCR sequences from scRNAseq data(*8*), we observed a highly variable clonal distribution of epitope-specific populations ranging from highly diverse to nearly monoclonal populations, which were found in HLA-B*07:01 GPR (CL-MCRL) and HLA-A*03:01 HYP (CH-3256) epitopes, respectively (Fig. 3F, table S6). We next quantified clonotype diversity by Shannon Evenness scores (SEv)) and found less diversity in responses with high magnitude of IFN-γ, also supported by a linear fit (R^2^=0.13, p=0.034) (Fig. 3G-H). These results suggest a divergent molecular signature and larger clonal expansion in cells associated with high IFN-γ responses.

### Distinct evolutionary trajectories of T_PEX_ in acute phase of infection explain functional heterogeneity

In chronic progressors T_EX_ and T_PEX_ subsets were the most abundant, particularly in the early phase of infection (Fig. 4A). To identify the static and dynamic components of the evolution of these subsets, partition-based graph abstraction (PAGA) was utilised to represent single-cell clusters as nodes and trajectories as edges (Fig. 4B). Clustering occurred according to the magnitude of IFN-γ production, epitope specificity, and clone size (Fig. 4C, fig. S3A). Four evolutionary trajectories were inferred from the PAGA structure using pseudotime analysis, which were grouped in two families based on their routes of origin (Fig. 4D) and were supported by RNA velocity inference, and by the increase in DPI of each sample time point along each trajectory (fig. S3). We then identified gene profiles that evolved over pseudotime. The first family of trajectories (Ch_T1, Ch_T2) comprised of cells from high-magnitude IFN-γ responses, which had larger clonal expansion (Fig. 4C) and targeted viral epitopes undergoing faster immune escape (Fig. 1G). The second family (Ch_T3, Ch_T4) comprised of cells from lower magnitude IFN-γ responses, formed more diverse clonal repertoires, and targeted epitopes undergoing slower or no immune escape. To quantify the evolutionary trajectories of exhausted and memory subtypes, the rate of growth of each subset was calculated along the pseudotime (Fig. 4E). All trajectories showed presence of T_PEX_ in early phase however family 1 high-IFN-γ trajectories showed faster growth rates for T_EX_ when compared to family 2. These trends were supported by higher MFI values for CD38 and CD95 and expression of genes such as *CD27*, *HLA-DR* and *LCK*, supporting increased TCR signalling and activation (Fig. 4F). In contrast, family 2 trajectories expressed higher amounts of *FOS, JUNB, IL7R, TCF7* since early phase which reflected increased prevalence of T_PEX_ and T_ML_ cells. We next characterised the transcriptional dynamics in each trajectory and found that family 1 trajectories carried a profile of early activation with both exhaustion and cytotoxicity profiles, with early upregulation of genes from glycosylation and fatty acid pathways, e.g., *STMN1* and *ECHS1* (Fig. 4F, table S8). In late phase memory profile dominated with expression of *FTH1*, *CXCR4*, *LTB*, *SELL* and *TCF7*, with expressions of *ZFP36* and *MALAT1*, which are known inflammatory regulators(*15, 16*), that were highly correlated with pseudotime (Fig. 4G). GSEA confirmed lack of AP-1 transcription factor (*FOS*, *JUN*, *JUNB*) enrichment in early phase of family 1 (fig. S3 C, D). By contrast, family 2 were enriched for cytotoxic associated genes (e.g., *GNLY*), had reduced signature of exhaustion, but increased expression of genes associated with controlling inflammation (e.g., *DUSP1*, *IFITM3*, *MALAT1* and *JUNB)* (fig. S3E-F).

**Fig. 4.**
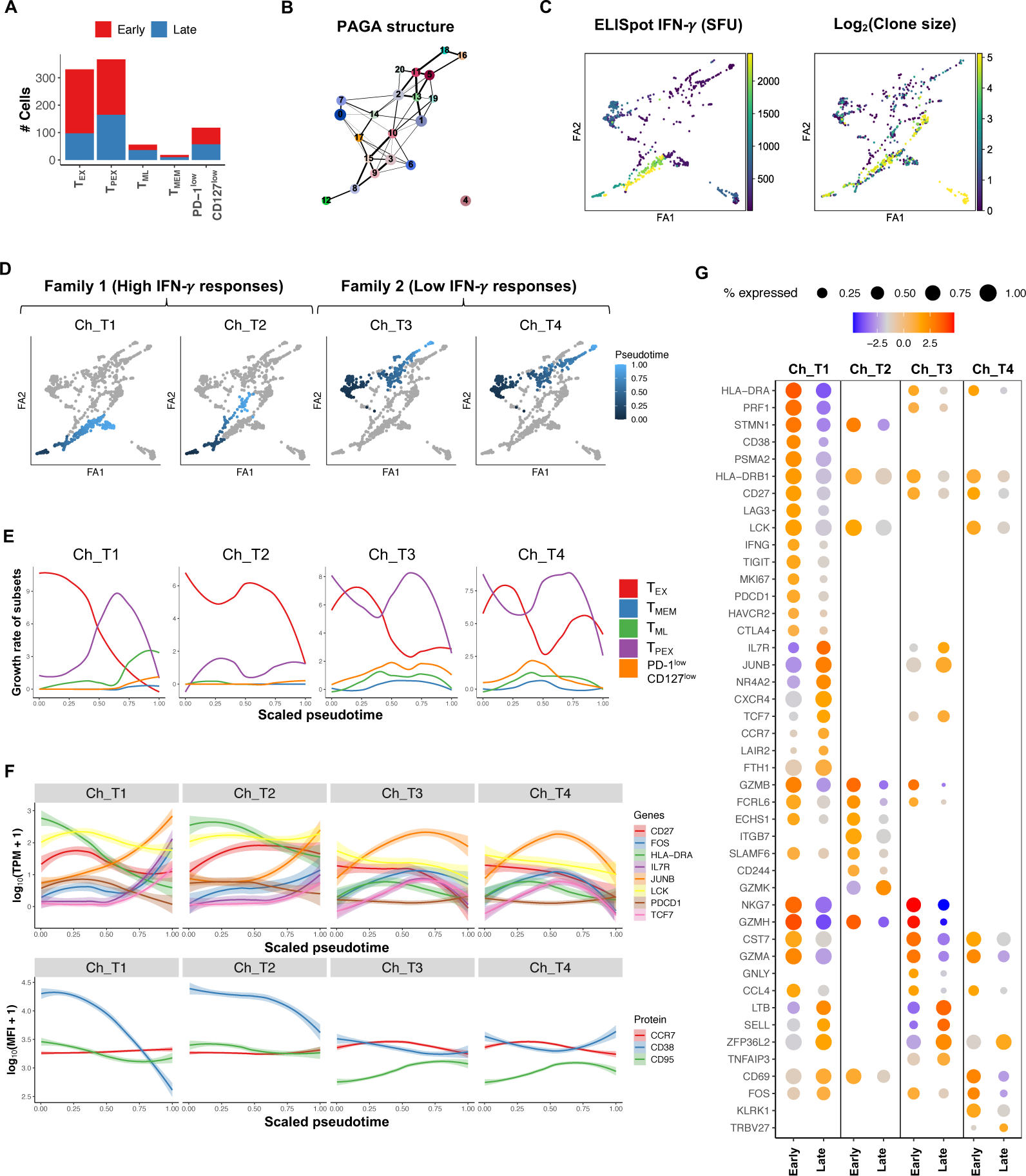
Distinct evolutionary trajectories of T_PEX_ in acute phase of chronic infection explain functional heterogeneity. **(A)** Distribution of cells from 7 chronic progressors by phase of infection (Early ≤ 120 DPI, late >120 DPI) in each phenotypic subset based on co-expression of CD127 and PD-1. **(B)** PAGA graphs of single cells from chronic progressors (n=903). The graph revealed both static and dynamic relationships by identifying clusters of cells as nodes and their connectivity (quantified by line weight as edges between clusters). **(C)** PAGA graphs (visualised using ForceAtlas2 layout algorithm (FA1, FA2)) coloured by IFN-γ (SFU/Million PBMC) and clone size. **(D)** Diffusion-pseudotime plots with colour gradients identifying four evolutionary trajectories grouped in two families: family 1 (Ch_T1 and Ch_T2_ and family 2 (Ch_T3 and Ch_T4). Trajectories are rooted in clusters from cells identified at the earliest sample time points. **(E)** Loess curves fitted to growth rates over pseudotime estimated from the calculated size of T-cell subsets, that are identified from the MFI values of PD-1 and CD127 for each cell. **(F)** Loess curves fitted to the expression of selected genes and surface proteins over pseudotime for each trajectory. **(G)** Dot plot of selected genes identified from differentially expressed genes between the early and late phases of each trajectory (p < 0.05, |log_2_(FC)| ≥ 1). The ball size represents the proportion of cells with non-zero expression. The colour represents fold-change relative to cells from the same trajectory at early or late phases. Early <=120 DPI, late > 120 DPI.

We performed a similar trajectory analysis to investigate the differentiation of 700 T cells from the two clearers, across three immune responses targeting two epitopes (fig. S4). These cells were mostly T_ML_, T_PEX_ and T_MEM_ cells and unlike those from chronic progressors, there were almost no T_EX_ cells (fig. S4A). We inferred four pseudotime trajectories (Cl_T1 to Cl_T4) rooted in early DPI clusters, and each with distinct phenotypic and gene signatures (fig. S5). Cl_T1 comprised GPR-specific T cells from viraemic and post-clearance samples (CL-3089) and developed from an early growing T_PEX_ population with an activation and cytotoxic (CD38^int^, *HLA-DRA*^+^, *NKG7*^+^, *PRF1*^+^) profile, which then persisted along with a T_ML_ subset (fig. S4E-G). The trajectories Cl_T2 and Cl_T3 shared the same root and consisted of an early T_MEM_ phenotype with low activation (CD38^low^), with T_ML_ cells dominating the late phase. These trajectories revealed high expression of AP-1 transcriptions factors, particularly during the early phase. Finally, Cl_T4, which included both GPR- and ATD-specific cells from CL-MCRL, comprised mostly of T_ML_ and T_PEX_ phenotypes, and revealed early expression of *FOS*, and dominant T_ML_ phenotype. GSEA confirmed these trends, particularly the higher activation and cytotoxicity of Cl_T1, and the distinct profile of Cl_T4 with reduced TCR signalling and cytotoxicity in early, and enrichment of oxidative phosphorylation in late phase (fig. S4I).

These findings collectively suggest that immune responses associated with chronic infection and fast rate of escape were associated with an early onset of T_EX_, and loss of expression of *TCF7* and AP-1 transcription factors expression, which contrasted with trajectories from viral clearance or slow rate of escape where T_PEX_ and T_ML_ dominated the early phase.

### Fate mapping reveals phenotypic plasticity within clonally expanded populations

We reasoned that analysing large clonal populations (Fig. 3H, table S6) would allow intra-clonal lineage tracing to demonstrate that single clones exhibit phenotypic plasticity through differentiation over the course of the infection. Over half of the cells formed expanded clones, and 63 (46%) clones persisted through both the early and late phases of infection, displaying a mix phenotype (Fig. 5A). The major phenotype in the early phase was T_EX_ whilst in the late phase it was T_PEX_.

**Fig. 5.**
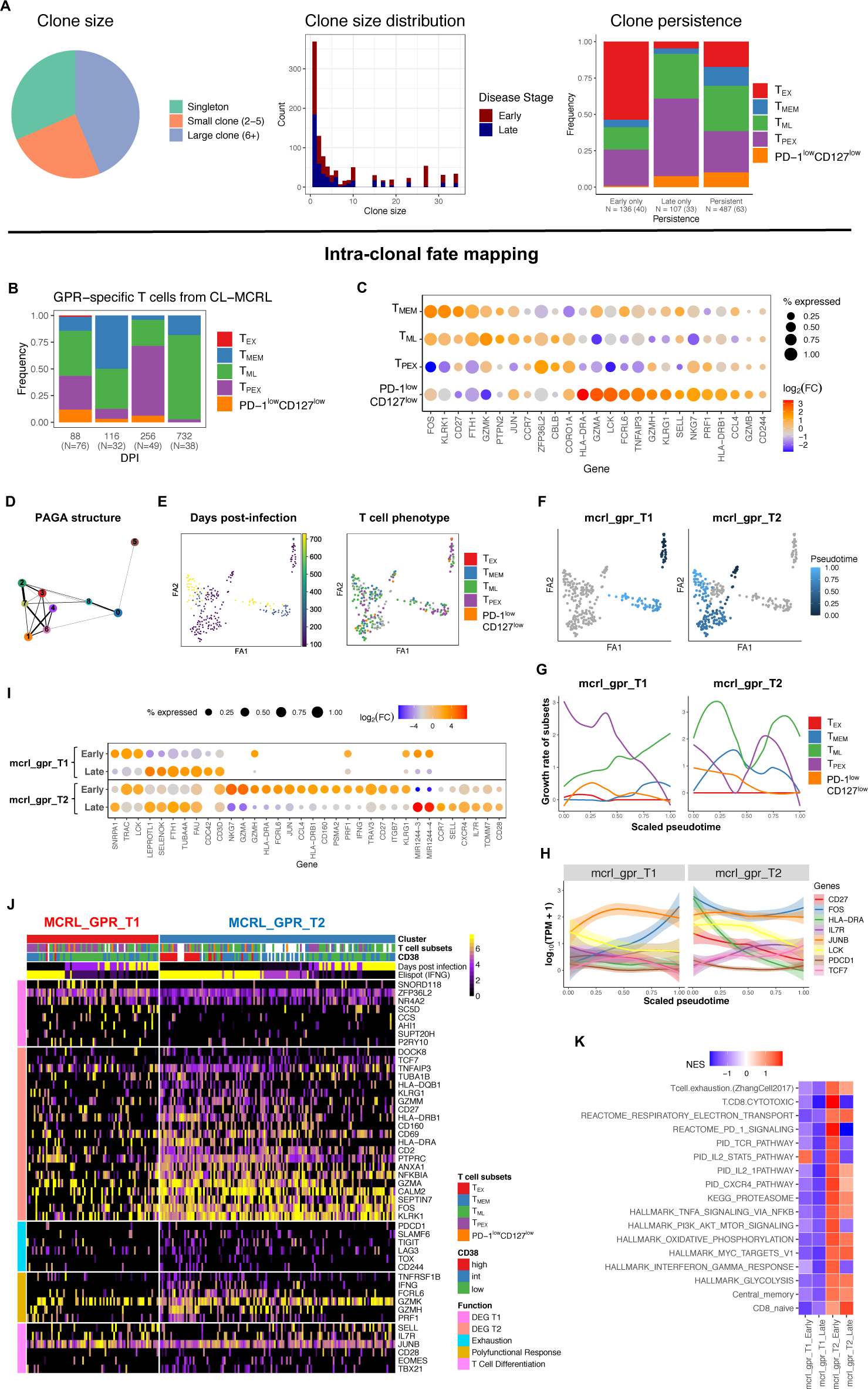
Fate mapping reveals phenotypic plasticity within clonally expanded populations. **(A)** Distribution of clones identified from the TCR sequences in each epitope specific response. Pie chart represents the frequency of clones grouped by size (number of cells with identical TCR*αβ* sequences). Histogram depicts distribution of clone size by disease stage (Excluding the large clone identified in CL-MCRL, see Fig. 5). Bar plot represents the distribution of clones by T-cell phenotype (TEX/exhausted: PD-1^high^CD127^low^, T_PEX_/progenitor-exhausted: PD-1^int^CD127^low^, T_ML_/memory-like: PD-1^high^CD127^high^, T_MEM_/memory: PD-1^low^CD127^high^). Shown are clones found only in early (≤120 DPI), only late (>120 DPI) phases, and observed at least once in both stages (persistent). Numbers represents cells and in brackets the unique number of clones. **(B)** Distribution of phenotype over time in cells identifying a monoclonal response (identical TCR*αβ* sequence) from the GPR-specific response in CL-MCRL (shown are 195 of the 244 cells available, due to unrecorded index sorting values). **(C)** Dot plot of selected genes from differentially analysis between phenotypic subsets of GPR-specific cells from CL-MCRL (p < 0.05) (A single cell had T_EX_ phenotype and was excluded from this analysis). **(D)** PAGA graph representation of GPR-specific cells (N=244) revealing 9 connected clusters (thickness of lines represent the probability of connectivity). **(E)** PAGA graphs (visualised using ForceAtlas2 layout algorithm (FA1, FA2)) coloured by the days post-infection of the samples and T-cell phenotype **(F)** Diffusion-pseudotime plots with colour gradients identifying two evolutionary trajectories. Trajectories are rooted in clusters from cells identified at the earliest sample time points. **(G)** Loess curves fitted to growth rates estimated from the calculated size of T-cell phenotypic subsets over pseudotime for each trajectory. Subsets are identified from the index sorted MFI values of PD-1 and CD127 for each cell. **(H)** Loess curves fitted to the expression of selected genes over pseudotime along each trajectory. **(I)** Dot plot of selected genes identified from differential expression analysis between early and late stages of each trajectory (p < 0.05, |log_2_(FC)| ≥ 0.9). Size of the ball represent the percentage of cells expressing each gene. **(J)** Heatmap of gene expression for differentially expressed genes between the 2 trajectories (mcrl_gpr_T1 and mcrl_gpr_T2). Genes were grouped by function and cells are annotated according to T-cell phenotype; legend “DEG T1” and “DEG T2” are representative genes identified from this analysis that are upregulated in mcrl_gpr_T1 and mcrl_gpr_T2 respectively. Shown on the top of the heatmap are the CD38 protein expression (MFI value per cell), days post-infection and magnitude of IFN-γ for each sample of origin. **(K)** Heatmap of enriched pathways identified from GSEA using differentially expressed genes between early and late phases within each trajectory. All pathways shown have p-values < 0.05 in at least one trajectory phase. NES: normalized enrichment score.

We considered the high IFN-γ monoclonal GPR-specific population in subject CL-MCRL to comprehensively track phenotypic plasticity (n=244 cells). This clone exhibited extraordinary levels of phenotypic plasticity- the T_PEX_ phenotype was present at all timepoints, and additionally major transitions were observed from T_PEX_ to memory (T_MEM_, T_ML_) phenotypes following viral clearance (Fig. 5B). T_PEX_ cells were enriched for *ZFP36L2*, *CORO1A*, *NKG7*, but had reduction in *FOS, JUN, LCK, GZMA,* and *FTH1*, suggesting an intermediate profile between memory and effector cells (CD127 ^low^ PD-1^low^) (Fig. 5C, table S8). The PAGA structure from these cells revealed clusters segregated by sample timepoint and phenotype (Fig. 5D-E). Two trajectories (mcrl_gpr_T1 and mcrl_gpr_T2) were inferred (Fig. 5F). Despite similar expression of TCR signalling genes, mcrl_gpr_T1 displayed an early T_ML_ phenotype, while mcrl_gpr_T2 displayed a rapid decline in the growth rate of T_PEX_ along the pseudotime, and a sustained accumulation of T_MEM_ and T_ML_ phenotypes (Fig. 5G). Notably, mcrl_gpr_T2 revealed activation (*CD27*, *HLA-DRA*, CD38^high^) and cytotoxic profile, including *IFNG* (Fig. 5H). Interestingly, the microRNA *MIR1244* reliably distinguished the two trajectories, being expressed early in mcrl_gpr_T1 and late in mcrl_gpr_T2. GSEA between the early and late phases of the trajectories revealed that mcrl_gpr_T2 had a significant increase in *IFNG* response, exhaustion and cytotoxicity during early phase, whilst metabolic associated pathways remained stable (Fig. 5J).

In summary, this analysis traced the evolution of multiple T-cell phenotypes within a single clone, demonstrating phenotypic plasticity and revealing distinct evolutionary trajectories corresponding to two lineages, one responsible for IFN-γ secretion and rapid loss of T_PEX_ and growth of T_ML_, while the other trajectory revealed a sustained T_PEX_ and T_M_.

### Epigenetic heterogeneity associated with responses targeting conserved and escaping epitopes

To further understand the molecular differences between responses targeting conserved or rapidly escaping epitopes, we performed bulk ATAC-seq to identify chromatin accessibility profiles on HCV-specific and total CD8^+^ T cells in the early phase of infection. Four responses were included in this analysis: the GPR-specific monoclonal response (CL-MCRL), two responses associated with rapid immune escape in chronic progressors (HPV and KLV epitopes), and the CIN-specific (CH-3023) with no escape identified. A principal component analysis (PCA) on accessible regions revealed clustering based on subject (Fig. 6A), and as expected, T cells from the monoclonal response were separated from the other populations derived from chronic progressors.

**Fig. 6.**
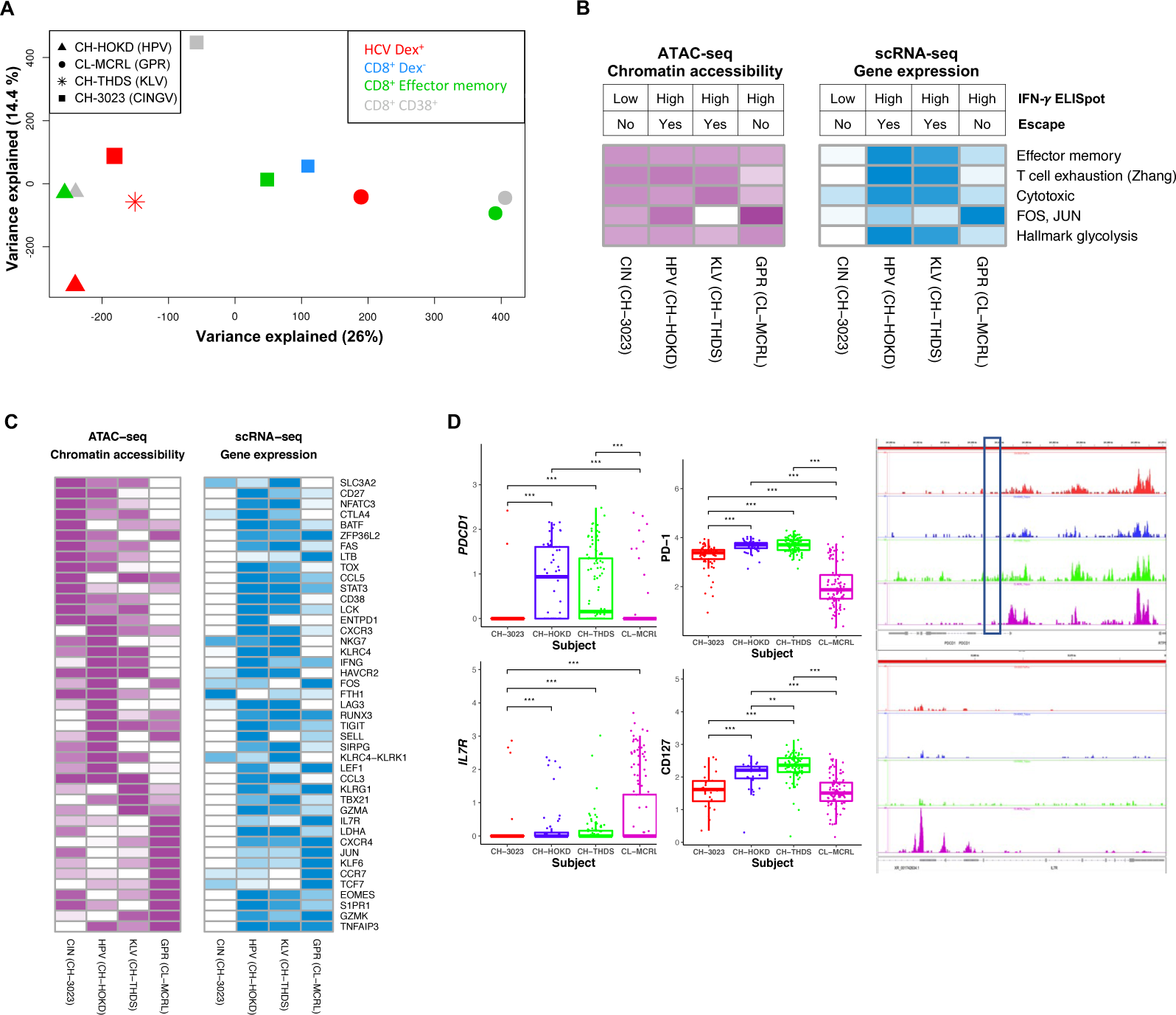
Epigenetic heterogeneity associated with responses targeting conserved and escaping epitopes. **(A)** Principal component analysis (PCA) from chromatin accessible sites of HCV-specific T-cell responses (Dextramer positive (Dex^+^) and subsets of CD8^+^ T cells, from samples within 120 days post-infection. **(B)** Heatmaps of average chromatin accessibility (in purple) and gene expression (in blue) for selected gene modules across four early epitope specific responses (DPI ≤ 120). Colour represents the average of the scaled genes in each gene set (Darker colours represent higher gene expression or chromatin accessibility). Responses are annotated by the corresponding IFN-γ ELISpot value and epitope escape. **(C)** Heatmaps of chromatin accessibility (in purple) and gene expression (log_10_(TPM+1)) (in blue) for selected genes, colours as in (B). **(D)** Comparison between gene expression (log_10_(TPM+1)) (left), protein expression (log_10_(MFI+1)) (middle) and chromatin accessibility (right) of selected genes for each of the 4 responses as per (B). Gene and protein values are from single cell data and from same sample time points as ATAC-seq data. Box highlights the intragenic *PDCD1* cis-element associated with terminal exhaustion^18^.

We then compared chromatin profiles of HCV-specific cells with the available single-cell gene and protein expressions measured at the same time points (Fig 6B-C). The responses in chronic progressors had increased chromatin accessibility at exhaustion (e.g., *ENTPD1* (encoding CD39)) and cytotoxic loci (*CCL3*, *KLRC4*-*KLRK1*), and consistent with gene expression was their decreased accessibility of *FOS, JUN* (Fig. 6D). Interestingly, the increased accessibility in exhaustion and effector memory genes in the CIN-specific response compared to the other responses, corresponded with a reduction in gene expression profiles. As this response targeted a conserved epitope, it is likely that lack of immune escape is associated with increased chromatin opening in these genes, as previously shown(*17*). As expected, *IL7R* accessibility was higher in CL-MCRL-GPR, while *PDCD1* locus showed increased opening in HPV- and GPR-specific responses. Notably, all populations lacked opening of the intragenic *PDCD1* cis-element associated with terminal exhaustion(*18*) which reinforced that the populations from this study are indeed precursors of terminal exhaustion (Fig. 6D).

Taken together, these molecular and epigenetic profiles demonstrate that in early phase of infection, T cells associated with early high functional responses have a distinct epigenetic control and early onset of both exhaustion and immune escape, while those in clearers revealed increased accessibility and expression of AP-1 transcription factors.

### Lineage tracing explains phenotypic and functional evolution of HCV-specific T cells

We summarised all the molecular and functional data from chronic progressors and clearers, and proposed a revised model of the role of CD8 T cells in controlling viral infection. While T_PEX_ cells were consistently found in trajectories from both disease outcomes and from early phase of infections, viral clearance was dominated by T_ML_, while chronic progressors revealed early onset of T_EX_ (Fig. 7A). Gene expression of cytotoxicity and activation were positively correlated with the magnitude of IFN-γ response, whilst memory markers and AP-1 transcription factors were negatively associated (Fig. 7B). The chronic trajectories revealed early expressions of exhaustion, effector, and metabolic functions proportionally to the rate of escape, while trajectories from clearers featured sustained memory markers and increased levels of *FOS* and *JUN* (Fig. 7C). To dissect the impact of reduced antigen stimulus in high IFN-γ responses, we compared the early phases of the Ch_T1 and mcrl_gpr_T2 trajectories (fig. S6). Despite the early presence of T_PEX_ and signature of polyfunctional response across these trajectories, Ch_T1 had increased activation, and exhaustion markers, while mcrl_gpr_T2 expressed more memory-like genes, regulatory genes (*ZFP36, DUSP1*, *DUSP2)*, and *FOS* and *JUNB.* In contrast, comparison between high and low magnitude chronic trajectories, revealed reduced exhaustion and cytotoxicity profiles in the latter, but increased memory and granzyme markers.

**Fig. 7.**
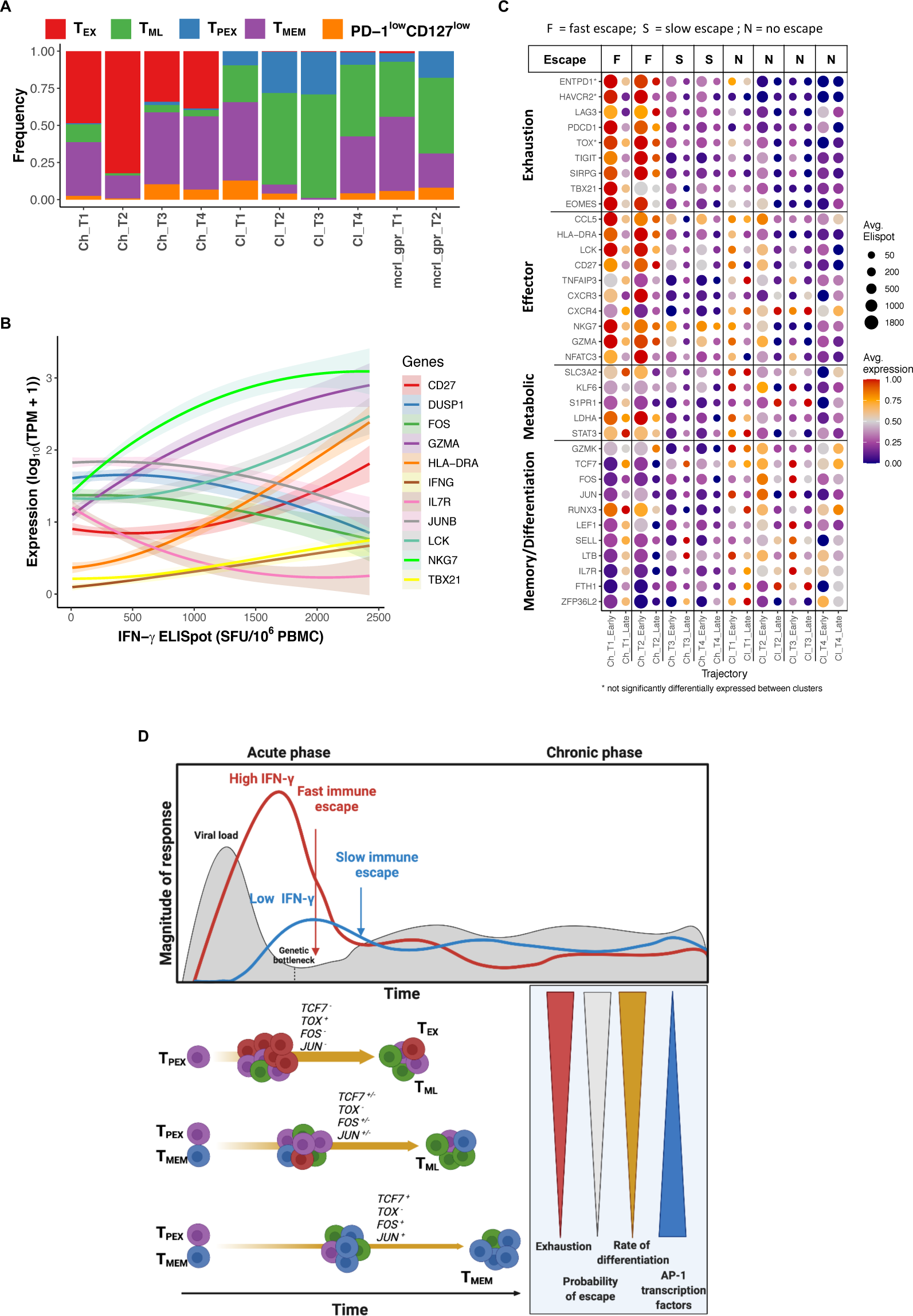
Lineage tracing explains phenotypic and functional evolution of HCV-specific T cells. **(A)** Distribution of T-cell phenotypes across the inferred evolutionary trajectories from chronic progressors, clearers and specifically for the intra-clonal analysis of the monoclonal response in CL-MCRL. **(B)** Loess curve fit of selected genes as a function of the magnitude of IFN-γ response. **(C)** Dot plot summarising the differences between early (<= 120 DPI) and late (>120 DPI) phase of chronic and clearer trajectories. Colour represents scaled averages of gene expression (log_10_(TPM+1)). Ball size represents average IFN-γ ELISpot (SFU/million cells). **(D)** Proposed model describing the differentiation and evolutionary dynamics of HCV-specific CD8^+^ T cells. The rate with which T cells undergo differentiation and clonal expansion determines the extent of exhaustion and the probability of viral immune escape.

Based on these findings, we developed a model of CD8^+^ T cell differentiation in early infection (Fig. 7D). In this model, it is assumed that the timing of clonal expansion varies between T cells due to stochasticity in antigen stimulus, TCR affinity and other factors. A strong stimulation early on promotes rapid clonal expansion and differentiation of T_PEX_ into T_EX_ and consequently high magnitude of IFN-γ response. This rapid response exerts an immunological pressure on viral evolution determining rapid immune escape. In contrast, delayed expansion results in a decreased probability of T_PEX_ differentiating into T_EX_ and favours memory precursors differentiating into T_ML_ and T_MEM_. Rapid expansion and differentiation of T_PEX_ are characterised by the loss of *TCF7* and AP-1 transcriptional factors (*FOS* and *JUN)*. In contrast, a slower differentiation leads to sustained AP-1 gene expression, dominating memory subsets, and lack of T_EX_. This model hence suggests that early rapid differentiation and expansion of antiviral cytotoxic T cells significantly contribute to viral evolution and early exhaustion.

## DISCUSSION

In this study, we analysed a unique collection of samples from subjects followed longitudinally from onset of acute HCV infection and implemented a multi-omics approach across virus and host to quantify the molecular, cellular and functional dynamics of epitope-specific CD8^+^ T cells. We found unexpected molecular and phenotypic heterogeneity of T cells at the divergence between chronic progression and acute resolution and discovered that the early clonal expansion and magnitude of IFN-γ response were predictive of viral immune escape and early onset of exhaustion. This contrasted with the sustained functional response observed in clearers, associated with a less differentiated memory-like phenotype, and expressing AP-1 transcription factors, *FOS* and *JUN.* These findings provide compelling evidence against the current model of a broad and high magnitude response predictive of viral clearance. Instead, optimal and sustained T-cell responses may be best achieved with moderate activation which avoids immune-driven selection of escape variants and early onset of exhaustion.

A key question in T-cell biology is how exhaustion is formed and then progresses from the progenitor cells to terminal status(*19*). The results of this study revealed that the early onset of exhaustion is a consequence of high magnitude, clonally expanded response, potentially representing a tolerance mechanism to control harmful cytotoxicity and excessive damage to the host(*20*). The rapid immune escape which results from early high magnitude IFN-γ responses leads to increased memory subsets as previously shown (*6*). These findings support a model whereby the maintenance of T-cell exhaustion in chronic phase is the result of prolonged responses against conserved epitopes targeted by low affinity T-cell repertoires. This conclusion is supported by the LCMV model where high-affinity T cells secrete high levels of IFN-γ against early dominant epitopes but are more extensively deleted compared to lower affinity T cells(*21*). It is conceivable that over the course of the infection new viral variants eliciting strong responses arise cyclically(*22*), and hence a similar dynamics that characterise early phase may be recurring in chronic phase. These conclusions will need validation to elucidate the dynamics of exhaustion and the role of escape variants in chronic infection.

Recent studies on HCV infection revealed that terminally exhausted but not progenitor exhausted cells are lost following clearance with antivirals or loss of antigen due to immune escape, and those that persist exhibit irreversible scars, both epigenetically(*23*) and functionally(*19, 24*) scars. Furthermore, transcriptional divergence has been identified as an early marker of cell exhaustion in HCV infection(*5, 25, 26*). Our study revealed complementary findings that suggest T_PEX_ can be found during early infection irrespective of outcome and that AP-1 transcription factors contribute to differentiation to a terminal exhaustion state. It is well known that these factors are involved in the regulation of T cell activation, and that cooperation with NFAT determine T cell anergy (*27*). *FOS* and *JUN* are downregulated in terminal exhausted cells(*28*) and play a role in induction of resistance to exhaustion in CAR T cells(*29*). Future studies are required to elucidate the exact molecular mechanism that control the expression of these transcription factors. We discovered that this early exhaustion phenotype evolved rapidly as a function of the rate of escape, and differed from that of terminal exhaustion, which has been previously associated with reduction of T-bet, and high levels of co-inhibitory markers(*9, 10*). Unexpectedly, a monoclonal response with high-IFN-γ response in a clearer provided an opportunity to further dissect the plasticity of progenitor cells and the existence of multiple fates committed to rapid generation of IFN-γ or long-term memory.

One of the limitations of this study was the absence of HCV-specific CD8 T-cell responses in the first 3 weeks post-infection, which are not observed in the blood(*30, 31*). During this phase, a broader range of T-cell repertoires may be recruited to the key site of viral replication in the liver, potentially with different affinities and breath of responses. Secondly, this study focussed on IFN-γ secretion for functional readout; however, antigen-induced production of other cytokines such as TNF-α, may reveal polyfunctional T-cell subsets, as it is also evident from our gene expression data.

Overall, this comprehensive multi-omics and longitudinal analysis of antiviral CD8 cytotoxic T-cell response in HCV provided new insights on how exhaustion arise and on the contribution of cytotoxic T cells to viral infection outcome. This knowledge is relevant for designing new vaccines and cellular therapies that elicit a successful and persisting CD8^+^ T-cell response that limit rapid onset of immune escape, in the context of both virus and cancer. The discovery of the important role of AP-1 transcription factors, should be further validated in other chronic infections and in the context of cancer therapies(*32*).

## MATERIALS AND METHODS

### Study Design

The study aimed to identify the molecular, cellular and phenotypic features of T cell responses in primary HCV infection. Subjects were selected from the Hepatitis C Incidence and Transmission Studies in prisons (HITS-p) and community (HITS-c) cohorts which recruited prospectively followed up high-risk injecting drug users from New South Wales, Australia on the basis of seronegative and HCV RNA-negative tests (*13, 33, 34*). All participants were consented from the University of New South Wales Human Research Ethics Committee (HC190074). Eligible participants had a lifetime history of injecting drug use and were documented to be anti-HCV and RNA-HCV negative in the 12 months prior to enrolment. Following initial detection of viremia, blood samples were collected frequently over a 24-week period until spontaneous clearance or chronic infection was established. Infection outcome was determined by considering whether each subject continued to test positive for viral RNA at six months following initial detection of viraemia. HCV antibody (Ab) and HCV RNA testing was performed as previously described (*35*).

### Subjects and samples

A total of 17 subjects were selected for this study. Fourteen subjects were analysed for viral sequencing, IFN-γ production, and flow cytometric phenotyping. Six subjects from these 14 were selected for scRNA-seq. An additional three subjects (CH-3221, CH-3132, CH-4059) were analysed by scRNA-seq only. All subjects selected for this study had longitudinally collected samples with estimated days post infection (DPI) previously described (*36*) and known infection outcomes. The estimated date of infection for early incident cases was estimated by subtracting the recognized mean pre-seroconversion window period of 51 days from the midpoint between the last HCV RNA^+^ HCV Ab^-^ time point and the first seropositive time point (*14*). For subjects with HCV RNA^+^ HCV Ab^+^ status at the initial infection time point, the estimated data post-infection was estimated as the mid-point between the last available HCV Ab^-^ and the first available HCV Ab^+^ timepoint (*36*). Blood samples were obtained longitudinally from participants enrolled in the HITS-p and HITS-c cohort. Blood samples were processed for isolation of PBMC using Ficoll-Paque gradients and then cryopreserved. Single cell multi-omics was performed using cryopreserved PBMC, which were thawed, quality checked for viable and live cells and then stained with dextramers for single cell sorting.

### Analysis of viral sequencing

Viral genomes were extracted from plasma and sequenced as described previously (*37*). Viral genomes for four subjects (CH-3023, CH-3240, CL-360, CL-MCRL) were sequenced and reported previously (*38*). Both consensus (Sanger) and deep sequencing were performed for 11 subjects, and in 3 subjects only consensus or partial genome sequences were available (table S2). T/F viruses were identified as previously described (*14*). Briefly, haplotype reconstruction was performed from the earliest available sample time-point from samples with 454 or Illumina sequencing to identify all peripheral circulating variants. Phylogenies were constructed containing all variants and for those with star-like structures, the central variant was identified as the T/F virus. PoissonFitter was applied to describe mutations arising from a single T/F virus under no selective pressure by the host immune response. For subjects with only Sanger sequencing, the consensus sequence at the earliest sample was identified as the T/F virus.

### Epitope selection

The set of HLA-restricted peptide epitopes tested in ELISPOT assays for each subject were determined from the circulating virus at the earliest viraemic timepoint. For each subject, this subset was selected from three major sets of epitopes. (i) set 1 consisted of autologous T/F virus or earliest viral consensus sequence HLA class I-restricted epitopes (9 to 11 amino acids) that were at least 90% homologous to previous experimentally confirmed epitopes from the Immune Epitope Data Base (IEDB [www.iedb.org]). If >1 T/F was present, then a selection was taken on epitopes that were predicted from the most dominant T/F (with the second virus having a frequency of occurrence <20%); (ii) set 2 consisted of epitopes identified by IEDB epitope prediction tools (using the IEDB-recommended procedure for 9 to 11 amino acids) from the earliest dominant viral sequence with a high predicted binding score (defined as a half-maximal inhibitory concentration [IC_50_] of 500 nmol), and 90% homologous to experimentally-confirmed epitopes available in the IEDB database; iii) set 3 consisted of potential immune escape variants identified from the longitudinal set of sequences associated with a nonsynonymous substitution that subsequently reached fixation over the course of the infection and with an increased IC50 of 500 nmol. As the total number of potential epitopes to be tested across these three sets was typically large (∼200-800 epitopes), a prioritised selection of up to 100 epitopes was undertaken for each subject to account for the total PBMCs available for the ELISPOT matrix testing approach and subsequent flow cytometric analyses (see below). A *post hoc* decision process was also undertaken in cases where the number of planned epitopes could not be tested due to limitations in the availability of viable PBMC following the thawing of frozen cells. In these cases, a subset of epitopes from the selected epitope pool was chosen by random sampling across the three pools. table S2 outlines the number of epitopes tested for each subject.

### Mathematical model to estimate the rate of immune escape

Viral epitopes were tested longitudinally and mutations occurring within the epitope regions were analysed using in-house scripts, which identified the mutations and their frequency of occurrence within the population. This tool takes in bam files generated from deep sequencing data, the reference sequence, the starting codon position and beginning and end position of the epitope region in nucleotide positioning based on the reference sequence as inputs. The escape variant was defined as any observed mutations away from the wild-type sequence occurring within the epitope. The frequency of the escape variant at each individual sample timepoint was estimated as the sum of the frequencies of each epitope variants carrying one or more mutations.

These longitudinal frequencies were then used to estimate the rate for CD8+ T-cell epitope escape by fitting the data to a population dynamics model that describes the dynamics of viral variants under the presence of cytotoxic T cell responses (*39*). The model predicts that the frequency of the escape variant *f(t)* is:

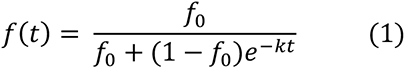

where *k is* the rate of escape. There is an assumption in this model that the escape variant is present during the initial phase of time (*t*) at zero, and its frequency is given by *f_0_*. In some cases where the escape variant was not observed in the earlier time points, an estimate of 1/(*n*+1) replaces a frequency of 0, where *n* is the average coverage of the corresponding time point from the deep sequencing data. This estimation was carried out in R using non-linear least-squares approach for non-linear models. The time needed for the escape variant to achieve 50% of the circulating viral population is obtained from the following formula:

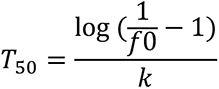

### IFN-γ ELISpot assay

ELISpot assays for IFN-γ production were performed as previously described (*34*). Briefly, samples were screened for responses in a matrix format using pools with ≤5 peptides. For each sample, groups of 150,000 PBMC were incubated overnight with peptides, a positive control, anti-CD3 antibody (Mabtech, Sweden), and three negative control wells with media only. Responses with greater than 20 SFU/million PBMC were considered positive. Positive responses were confirmed by stimulation with a single peptide and 200,000 PBMC. Positive responses were defined as exceeding the background level, defined as the mean plus three times the standard deviation of SFU in the negative control wells.

### Flow cytometry

Peripheral blood mononuclear cells (PBMCs) were thawed in RPMI and washed with PBS containing 1% BSA. Cells were stained with PE-conjugated HCV-specific class I dextramers (Immudex, Copenhagen, Denmark) at room temperature, followed by viability staining (LIVE/DEAD^TM^ fixable blue for analysis or fixable yellow for sorting) (Invitrogen, Carlsbad, CA) and staining with panels of surface or intracellular antibodies detailed below.

Flow cytometry was performed using the LSR Fortessa analyser, and FACSAria III and Influx sorters (BD Biosciences, San Diego, CA). Flow cytometry data was analysed using FlowJo version 10.1 (BD Biosciences, San Diego, CA). Gates for CD127, PD-1, CD38, and Tim-3 were determined with fluorescence minus one (FMO) controls.

Antibodies were sourced from BD Biosciences unless stated otherwise. Two panels of antibodies were used for flow cytometry analysis. The first surface marker panel included: BV421 and PerCP/Cy5.5 anti-TIM-3 (clones B8.2C12 and RMT3-23, BioLegend), BV510 anti-PD-1 (EH12.1), BV605 anti-CD38 (HB7), BV650 anti-CD127 (HIL-7R-M21), FITC anti-CD4 (RPA-T4), PE-Cy5 anti-CD19 (HIB19), PE-Vio770 anti-2B4 (REA112, Miltenyi Biotec), AF647 anti-CD160 (BY55, BioLegend), AF700 anti-CD8 (RPA-T8), APC-Cy7 anti-CD3 (SK7), BV650 CD127 (HIL-7R-M21), BV421 CCR7 (150503) PE-Cy-7 CD45RO (UCHL1). Dextramers (Immudex) were conjugated with PE. For KLRG1 staining, cells were incubated with primary antibody Biotin anti-KLRG1 (2F1) followed by incubation with secondary antibody PE-CF594 Streptavidin. In a second panel, for the intracellular staining, cells were fixed and permeabilized using fix/perm buffer from transcription factor buffer set kit (BD Biosciences) at 4°C for 35 minutes and stained with against intracellular markers anti T-bet BV711 (clone O4-46), anti EOMES eFluor660 (WD1928, eBioscience) at 4°C for 30 minutes. The cells were then washed twice with Perm/Wash buffer (BD Biosciences) and fixed with PBS containing 1% paraformaldehyde.

A separate panel was used for flow cytometry for sorting and single-cell RNA-sequencing including: BV421 anti-CCR7 (clone 150503), BV480 anti-CD3 (UCHT1), BV650 anti-CD122 (MiK-β3), BV786 anti-CD95 (DX2), APC anti-CD38 (HB7), APC-R700 anti-CD8 (RPA-T8), PD-CF594 anti-PD-1 (EH12.1), PE-Cy5 anti-CD19 (HIB19), PE-Cy7 anti-CD127 (HIL-7R-M21), FITC anti-CD45RA (HI100), PerCP/Cy5.5 anti-KLRG1 (SA231A2, BioLegend). Dextramers (Immudex) were conjugated with PE.

### ATAC-seq

ATAC-seq was performed using samples from four subjects at the following timepoints: CH-HOKD 96 DPI, CL-MCRL 115 DPI, CH-3023 73 DPI, and CH-THDS 85 DPI. All timepoints were within the initial 120 DPI and up to 10000 cells were sorted from a combination of dextramer-positive CD8^+^, dextramer-negative CD8^+^, total effector CD38^+^CD8^+^, and total effector memory CD8^+^CCR7^-^CD45RA^-^ populations. For HCV-specific (dextramer) responses in subject CH-THDS (KLV-specific) two samples were utilised, and only one for the remaining responses. All HCV-negative responses were in duplicates or triplicates. A total of 16 libraries were generated. DNA library preparations were carried out using a previously published protocol (*40*). Briefly, target populations were sorted by flow cytometry (FACSAria III). Cells were then lysed and fragmented in a single reaction (12.5 µl 2X Illumina TDE buffer, 2.5 µl 1% Tween-20, 2.5 µl 0.2% Digitonin, 5 µl water, 2.5 µl Illumina TDE1 enzyme). Samples were incubated at 37°C for 60 minutes and purified using the Zymo DNA Clean and Concentrator-5 Kit (Zymo). Fragments were amplified according to the previously published protocol. The total number of cycles was selected to be the number of cycles required to reach one quarter of maximum qPCR fluorescence. Products were purified using Ampure XP beads (1.5X) and quality controlled using a TapeStation (Agilent Technologies). Libraries were sequenced by paired end (2x75 bp) to a depth of 25 million reads per sample (Illumina Miseq v3).

Sequenced reads were trimmed with Trim_Galore (version 0.4.5_dev) and aligned to GRCh38.p12 using BWA-MEM v0.7.17 (*40*) in paired-end mode with default parameters. The resulting bam files were deduplicated with MarkDuplicates (Picard) v2.19.0. Prior to further analysis, sample quality was verified using the ATACseqQC package v1.10.1 (*41*). Samples were normalised using the normOffsets() function, peak regions across samples defined by mergeWindows() with tol=1000L and max.width = 5000L, and regions annotated by detailRanges() from the Bioconductor package csaw v1.20.0 (*42*), with default values used unless otherwise specified.

### Statistical analysis

Statistical analyses were performed using GraphPad Prism 7.0 (GraphPad Software, Inc., La Jolla, USA) and R (*43*). Data was expressed as the median with interquartile range and analysed using non-parametric statistics. Analysis of ELISPOT data was performed assuming positive responses only with values above 30 SFU/million PBMC. Epitope-specific immune responses with a single value <30 SFU were excluded from downstream analysis. For flow cytometric data, comparisons of gated populations between infection outcome and between time windows were performed using Wilcox sum-rank tests, and p-values less than 0.05 were considered significant. Regression and correlation analyses were performed in R using function “lm” and “ggplots2” libraries. Regression analysis between magnitude of IFN-γ production and rate of escape, the maximum value of the IFN-γ ELISPOT measured for each epitope within the first 16 weeks post-infections was considered.

## Acknowledgements

The HITS-p and HITS-c investigators include Andrew Lloyd, Lisa Maher, Kate Dolan, Paul Haber, William Rawlinson, Carla Treloar and Greg Dore. Flow cytometry and sequencing were supported by staff at the UNSW Flow Cytometry and Ramaciotti Centre core facilities. This research was supported from National Health and Medical Research Council of Australia (NHMRC) - Project Nos. APP1121643, 1027551, 1060199, Partnership No. 1016351, and Program Nos. 510488 and 1053206. The HITS-c cohort was supported by the UNSW Hepatitis C Vaccine Initiative and NHMRC Project Grant No. 630483. F.L., A.R.L. and R.A.B. are supported by NHMRC Research Fellowships (Numbers: 1128416, 1041897 and 1084706). M.R.P., P.L., and C.C. were supported by an Australian Government Research Training Program (RTP) Scholarship. S.G. was supported by the NHMRC grant APP1148284. K.K. is supported by the NHMRC grant APP1148284. Leadership Investigator Grant (#1173871), T.H.O.N. is supported by NHMRC Emerging Leadership Level 1 Investigator Grant (#1194036).

## Author contributions

F.L. designed and led the study with support from R.A.B. and A.R.L. A.R.L. designed and led the cohort studies. M.R.P. and E.K. performed the ELISpot experiments. C.C., M.R.P. and E.K. performed the flow cytometry experiments. C.C., M.R.P., F.L. and E.K. analysed the ELISpot and flow cytometry data. S.G. and A.R.L. provided support for the ELISpot, flow and viral epitopes identification analyses. C.C., T.N.A. and A.E. performed the scRNA-seq experiments. S.G., A.R.L., K.K. and T.H.O.N. contributed reagents. J.S., B.H., J.L.P., R.H.L., M.G., C.C., T.P., and W.v.d.B. analysed the scRNA-seq data. T.N.A. and C.C. performed the ATAC-seq experiments. J.S. and T.P. analysed the ATAC-seq data. F.L., J.S. and C.C. performed remaining analyses and generated figures. F.L., C.C. and J.S. wrote the manuscript. All authors read the manuscript and provided comments.

## Data and materials availability

Raw and processed data for this study are available at ArrayExpress (accession numbers: E-MTAB-XXX for scRNAseq, E-MTAB-XXX for ATAC-seq). Viral deep sequencing data are all available upon request to the authors. All other data needed to evaluate the conclusions are present in the paper or the Supplementary Materials.

## Supplementary Materials

### Material and Methods

#### scRNA-seq library preparation

Index sorted cells were collected into 96-well plates. Single-cell RNA-seq libraries were generated with a modified SmartSeq2 protocol as previously described (*1*). Briefly, a modified protocol from Picelli et al (*2*) was used to reduce volumes and concentrations of reagents. Sequencing of the libraries was performed on the Illumina Next-Seq or Mi-Seq machines with high throughput kit 150bp and 250 bp respectively.

#### scRNA-seq bioinformatic analysis

The overall bioinformatics pipeline for the analysis of the scRNA-seq data was conducted mostly as previously reported by our group (*3*) with some modifications as outlined below.

#### Sequence Analysis

Raw sequencing reads were trimmed using Trimmomatic (v0.39) (*4*) and aligned to the reference genome GRCh38.p13 using STAR (v2.7.3a) (*5*). Gene expression was quantified using RSEM (v1.3.1) (*6*) and Ensembl gene annotation release 99. Scaled transcripts per million (scaled TPMs – TPMs scaled so that the total count per cell is equal to the total aligned read count) were extracted from the RSEM results and used for further analysis.

#### Quality Control

Downstream analysis was performed in R using packages downloaded from Bioconductor 3.10. Cells were removed from each batch if they did not meet these criteria: less than 30% reads aligned to mitochondrial genes, total reads more than 30,000 and less than the 98% quantile for the batch, number of detected genes more than 400 and less than the 98% quantile for the batch, and number of detected genes less than the 25% quantile for detected genes in mini-bulk controls samples (of 30-50 cells) for the batch. The bulk samples were also removed at this stage. To assess the factors (sampling time point, subject, epitope, library size) contributing to the transcriptional variance between cells the function *getVarianceExplained* and *plotExplanatoryVariables* functions were used from the scater package in R. Genes expressed in less than 1% of cells were removed from the dataset. The top 7,500 variable genes were detected using the *FindVariableFeatures* method in Seurat with the “vst” option.

#### Clustering

The variance stabilising transform (VST) as implemented in Seurat (v3.2.2) (*7*) was used for normalisation and subsequent dimensionality reduction with UMAP and clustering. Principle components analysis (PCA) was performed on the VST normalised data using the 3,000 most variable genes. The first 30 principal components were selected as the most significant based on an elbow plot and used as input for UMAP and clustering. Clustering was performed using the shared nearest neighbour (SNN) modularity optimization-based clustering algorithm (*FindClusters*(resolution = 1, algorithm = ‘louvain’)) as implemented in Seurat, to obtain 13 clusters. Analysis of reference-based clustering was performed using SingleR (v1.0.6) (*8*).

#### Differential gene expression, trajectory inference and gene signature scores

Differential expression was performed with MAST (v.1.16.0) (*9*), using Wald tests to estimate p-values, with a p-value cut-off of 0.05. Differential expression output across all the analyses is reported in table S8. Normalised gene counts using the deconvolution method (*10*) was utilised for differential expression analysis.

PAGA analysis was performed through SCANPY (v1.7.1) with parameters as recommended (*11*). Log transformed scaledTPM matrix was used for initial pre-processing steps and visualization using sc.pp.neighbors and a coarse-grained and simplified graph using sc.tl.paga(). Clusters were calculated using sc.tl.louvain() and visualization was performed using sc.pl.paga() and sc.pl.draw_graph(). Diffusion pseudotime was inferred using sc.tl.dpt() by manually assigning an initial iroot value. Scaled pseudotime were used with LOESS smoothing (Fig 4e-f, Fig 5g-h, Extended data Fig 4f-g) and were calculated as uniformly distributed mapping of the diffusion pseudotime inferred from sc.tl.dpt(). This transformation preserves the cell order and serves to reduce the effect of gaps in pseudotime on the LOESS smoothing. Phenotype growth rates (Fig 4e, Fig 5g, Extended data Fig 4f) were calculated as the difference in the number of cells for each phenotype across scaled pseudotime windows of size 0.05. These values were then plotted using the *geom_smooth()* R function with default parameters.

Signature scores were computed from the single cell transcriptomic matrix as the average log(TPM+1) of all genes in the signature. Gene signatures were taken from MSigDB, while for functionally distinct T cell subsets signatures were obtained from published data (*12, 13*).

#### Gene set enrichment analysis (GSEA)

Gene set enrichment analysis was performed using the R package fsgea (v1.16.0). Normalised enrichment scores (NES) were assessed using the fgsea(…, maxSize = 500, nperm = 10000) function across the curated Molecular Signatures Database (MSigDB) Hallmark, C2 curated gene sets comprising REACTOME, KEGG and Canonical gene sets PID), C5 Gene ontology, and C7 Immunological signature. Customised gene signatures for T cell phenotypes are reported in table S9.

GSEA was performed for each evolutionary trajectory to identify enriched gene sets in the early and late phases (Figure 4, and 5). In a second application (Figure 6, and 7) GSEA was applied to identify enriched gene sets in each trajectory phase (split by early or late) via comparison with all other trajectories.

#### RNA velocity

Unspliced and spliced transcripts were generated from each cell’s BAM file using velocyto (v0.17.17) (*14*), and were analysed using scVelo (v0.2.2) (*15*) to derive counts and to estimate velocity parameters. Velocities were then visualised onto the PAGA embedding. The velocities were estimated using the stochastic model (sc.tl.velocity() and sc.tl.velocity_graph()).

#### TCR repertoire analysis

TCR full length alpha and beta sequences were obtained from scRNA-seq data using the software tool VDJPuzzle(*1*). In two subjects, single cell TCR Sanger sequencing were also generated, utilising established protocol (*16*). For subject CH-3023 Sanger sequencing data were generated for the same two epitope-specific responses with available scRNA-seq, at 75, 101 (additional timepoints), and 196 DPI (existing timepoint). For subject CH-240, all TCRs were generated using only Sanger sequencing from two epitope-specific responses at 71, 99, and 140 DPI (additional timepoints) (see table S7). For scRNA-seq data, the α and β chains with the highest expression were used where multiple chains were reported in a single cell. Clones were defined as cells with identical TCR CDR3 amino acid sequences in both α and β chains. Shannon entropy (SE) was calculated using the *entropy* function (with parameter “.base = exp(1)”) in the Immunarch R package (v0.6.5), utilising the CDR3 amino acid sequences of both α and β chains. Shannon evenness (SEv) was calculated as SE/log(*N*) where *N* is the number of sequences used to calculate SE.

#### Codes

Customised in house scripts for analysis of viral genomes, functional and transcriptomics data are available upon request.

## Supplementary Figures

**Fig. S1.**
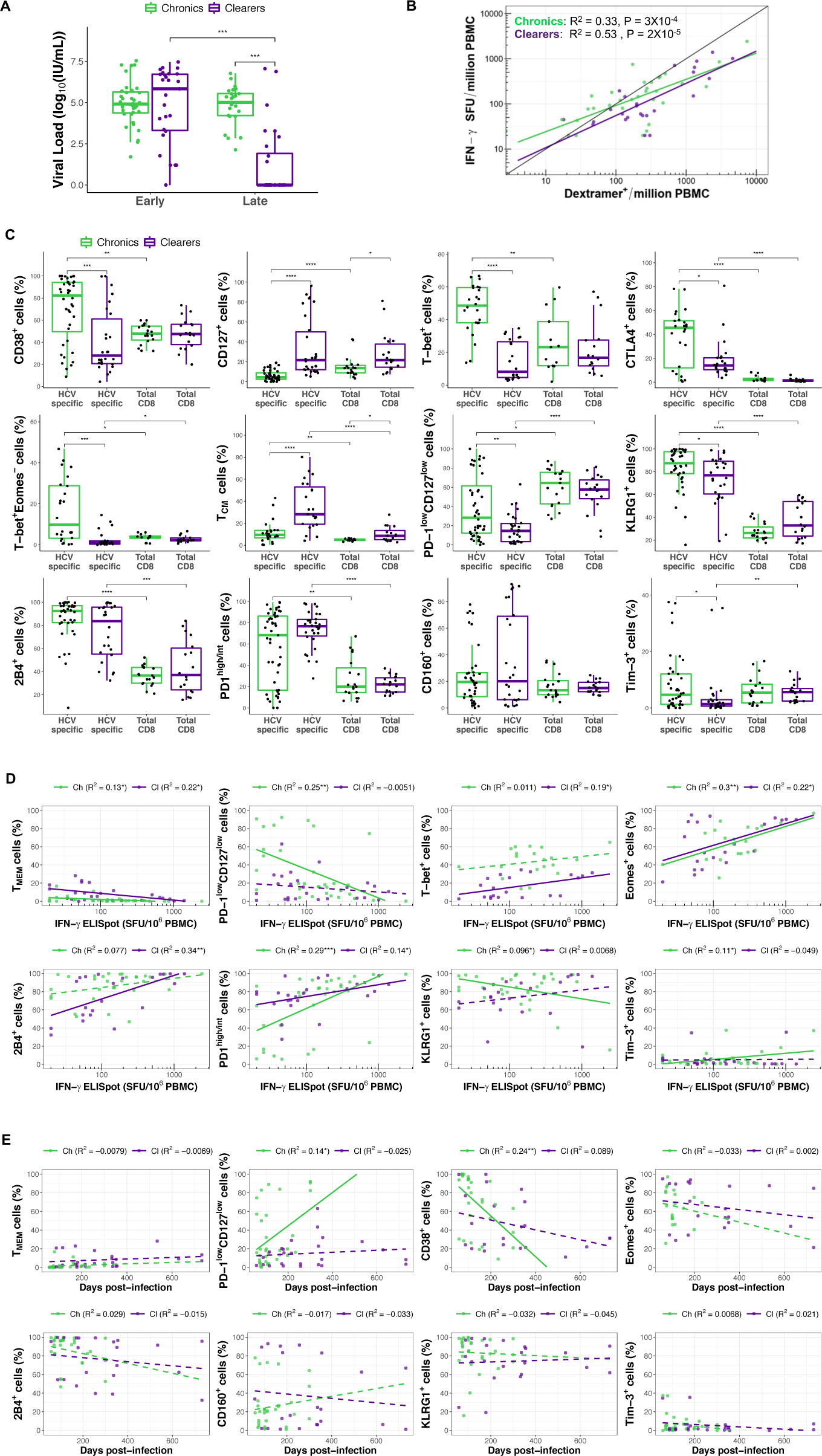
Phenotypic and functional characterization of HCV-specific CD8^+^ T cells. **(A)** Comparison of viral load values by phase of infection (Early <=120 DPI, late =>120 DPI) and disease outcome. **(B)** Scatter plot showing the correlation between IFN-γ ELISpot values (number of SFU per million PBMC) and number of epitope-specific CD8^+^ T cells (number of dextramer positive CD8^+^ T cells per million PBMC). Linear regressions were obtained for both disease outcome. The black line represents the bisectrix, indicating a 1:1 correlation between both axes. **(C)** Box plots of the proportion of cells from each population with positive expression for markers measured by flow cytometry compared between HCV-specific CD8^+^ T cells in both disease outcomes, as well as total CD8^+^ T cells. Individual points represent populations from each subject’s sample time points, and from each epitope specificity. TCM: central memory. **(D)** Scatter plots showing the relation between the proportion of dextramer positive CD8^+^ T-cell populations with positive expression of phenotypic markers (measured by flow cytometry) and IFN-γ ELISpot values (number of SFU per million PBMC). Lines represent linear regression, adjusted coefficients and p-values are shown in the legend of each plot. Lines are dashed if insignificant (p-value > 0.05). T_MEM_/memory: PD-1^low^CD127^high^. **(E)** Same as (D) but proportions are plotted and regressed against days post-infection (DPI). Statistical comparisons between groups were performed with Wilcoxon Rank Sum test (* p < 0.05, ** p < 0.01, *** p < 0.001, **** p < 0.0001).

**Fig. S2.**
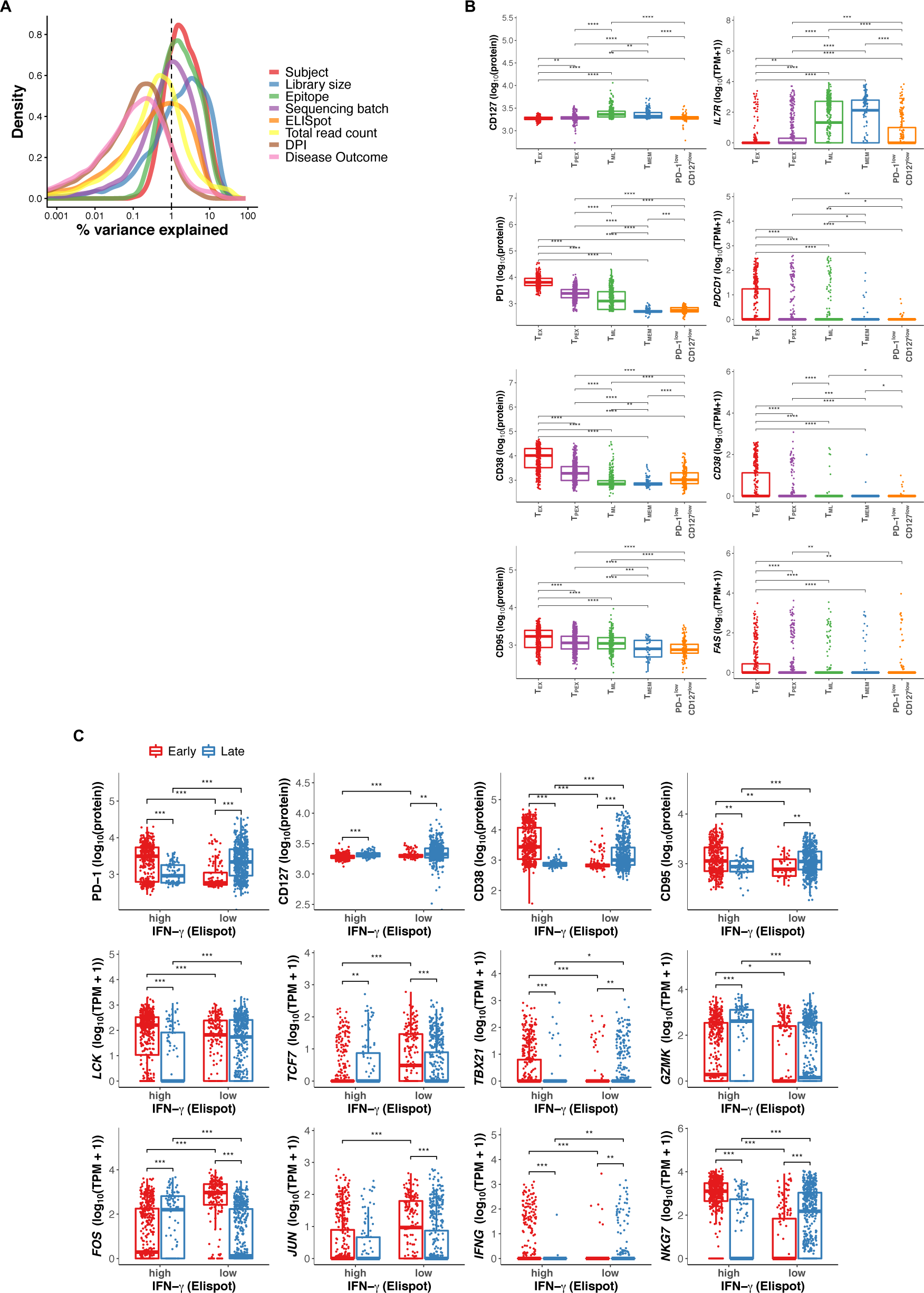
Molecular and phenotypic heterogeneity of HCV-specific CD8^+^ T cells. **(A)** Analysis of factors contributing to variability of gene expression in single cell data, including batch effects. Shown is the density plot of the percentage of variance explained. Each curve corresponds to one factor. **(B)** Comparison (box plots) of proteins (log_10_(scaled MFI + 1)) and corresponding genes (log_10_(TPM + 1)) expression levels between cells grouped by their phenotypes. **(C)** Similar to (B), comparison between cells grouped by disease stage (Early <= 120 DPI and late > 120 DPI) and by corresponding magnitude of response (High >211 (SFU), low <= 211 SFU). Statistical comparisons between groups were performed with Wilcoxon Rank Sum test (* p < 0.05, ** p < 0.01, *** p < 0.001, **** p < 0.0001).1

**Fig. S3.**
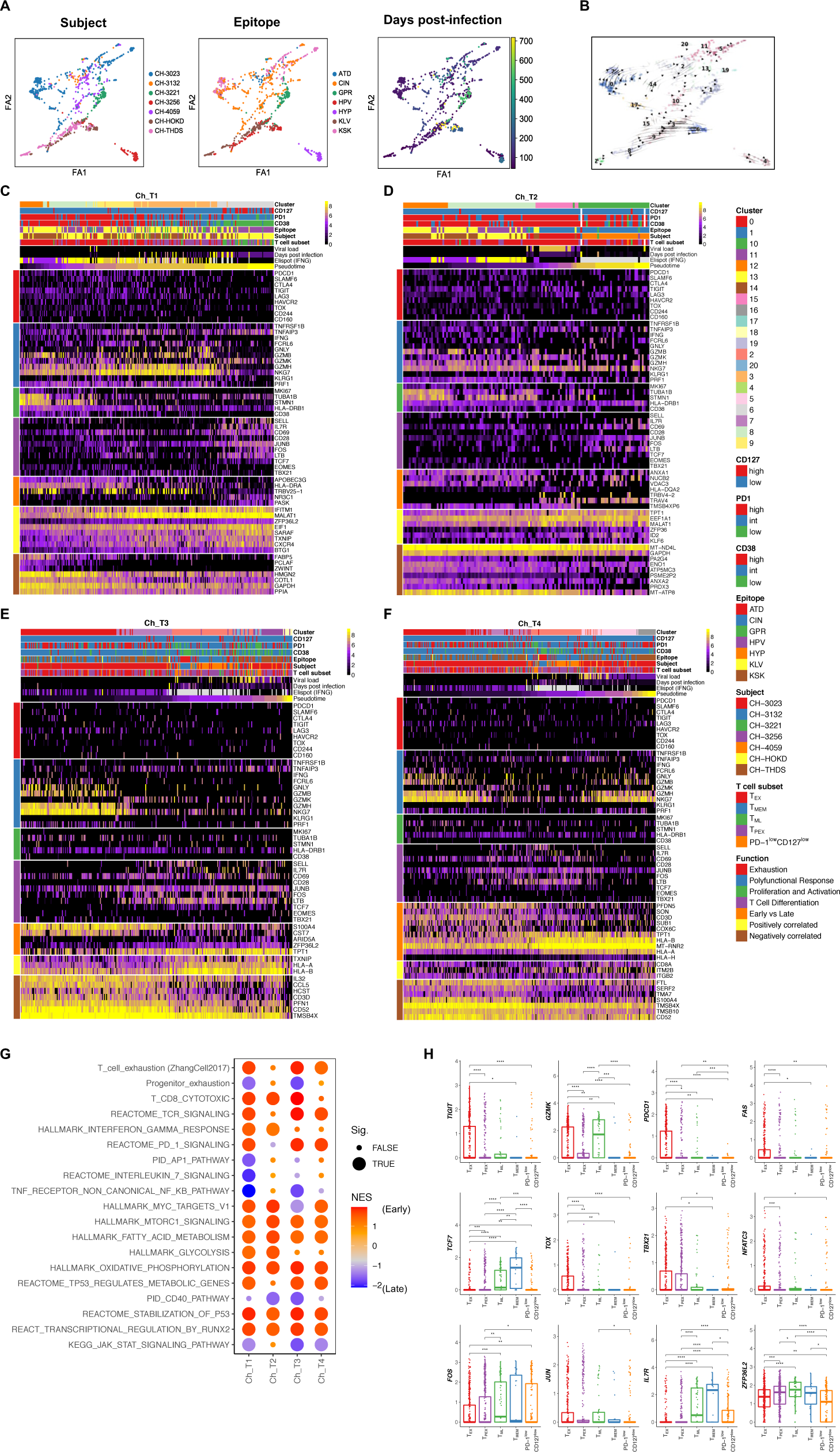
Trajectory analysis reveals molecular and phenotypic evolution of HCV-specific T cells in chronic progressors. **(A)** PAGA graphs (visualised using ForceAtlas2 layout algorithm (FA1, FA2)) from scRNA-seq data as per (Fig. 4B). Panels show cells coloured with different metadata features. (B) RNA velocities visualised on the PAGA graph revealed evolutionary trajectories consistent with those inferred using diffusion pseudotime. **(C-F)** Heatmaps of representative genes from cells forming family 1 trajectories (see Fig. 4), Ch_T1 **(C)** and Ch_T2 **(D)**, and family 2, Ch_T3 **(E)** and Ch_T4 **(F)** trajectories. On the top of each heatmap T-cell phenotype (index sorting), epitope, subject, viral load and IFN-γ ELISpot measured at the sample of origin are shown for each cell. Selected genes from differential expression analysis between early and late phase of infection, as well as genes correlated with pseudotime are shown. Genes were grouped based on their functions. Cells are ordered by pseudotime revealing positive correlation with the sampling time point (Days post infection). **(G)** Dot plot of enrichment scores from GSEA using differentially expressed genes between early and late stages within each trajectory independently. Normalised enrichment scores (NES) were calculated independently within each trajectory. Positive NES indicates enrichment of gene set early in the trajectory, and negative NES indicates late enrichment. Results were considered significant (Sig.) if p < 0.05. **(H)** Box plots of mRNA expression levels (log_10_(TPM + 1)) for selected genes in cells from chronic progressors defined by protein co-expression of PD-1 and CD127 (TEX/exhausted: PD-1^high^CD127^low^, T_PEX_/progenitor-exhausted: PD-1^int^CD127^low^, T_ML_/memory-like: PD-1^high^CD127^high^, T_MEM_/memory: PD-1^low^CD127^high^). Statistical comparisons between groups were performed with Wilcoxon Rank Sum test (* p < 0.05, ** p < 0.01, *** p < 0.001, **** p < 0.0001).

**Fig. S4.**
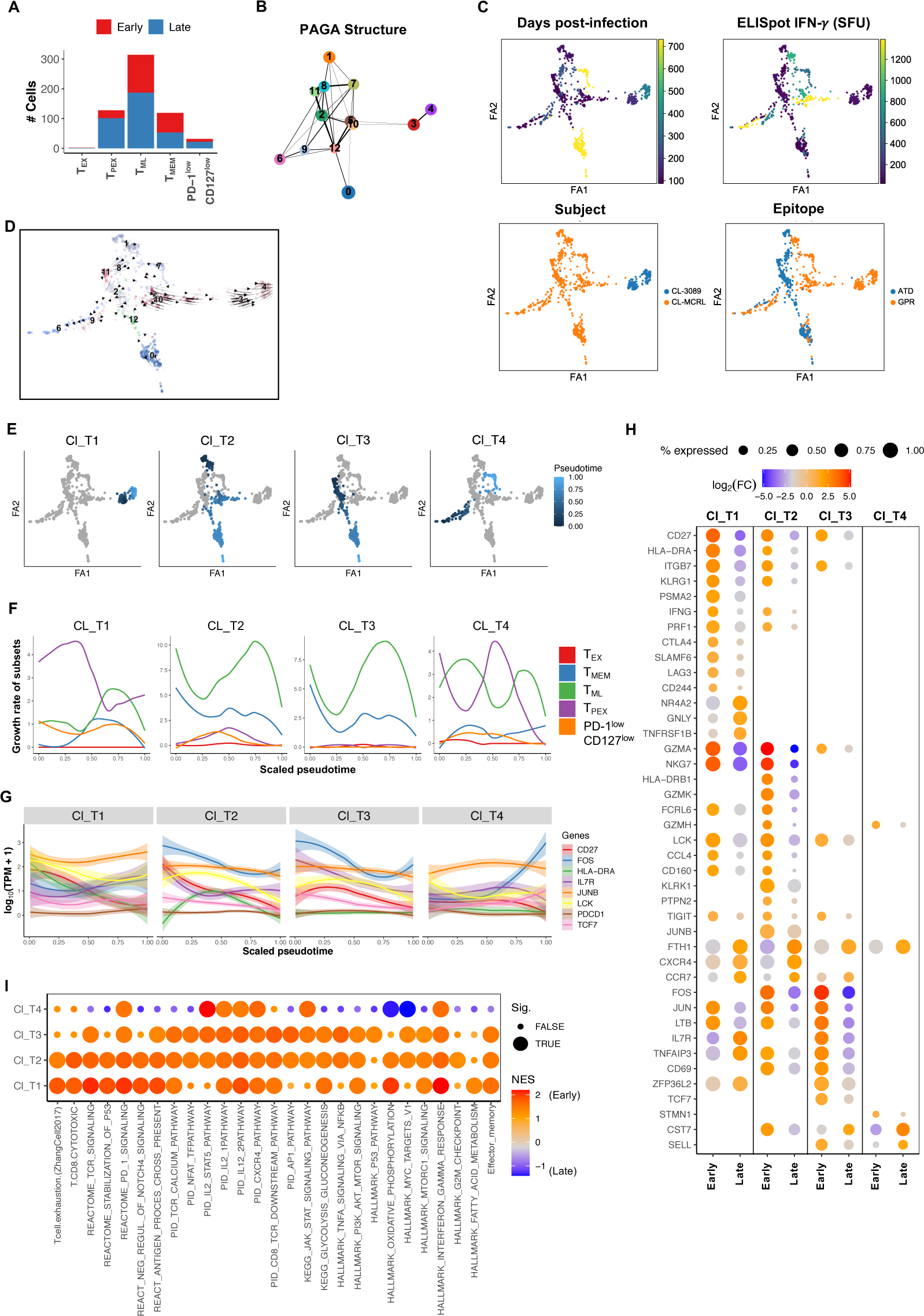
Trajectory analysis explains sustained IFN-γ production and evolution from T_PEX_ to memory-like phenotypes in clearers. Distribution of cells by phase of infection (Early ≤ 120 DPI, late >120 DPI) in each phenotypic subset based on co-expression of CD127 and PD-1. **(B)** PAGA graph of single cells from 2 clearers (n=700 cells) identifying clusters of cells as nodes and connectivity, quantified by line weight, as edges between clusters. **(C)** PAGA graphs (visualised using ForceAtlas2 layout algorithm (FA1, FA2)) coloured by infection features as per legend. **(D)** RNA velocities visualised on the PAGA representation. **(E)** Diffusion-pseudotime plots with colour gradients identifying the four selected trajectories. Ch_T1 and Ch_T2 were classified as family 1; Ch_T3 and Ch_T4 were classified as family 2. **(F)** Loess curves fitted to growth rates each phenotype from (A) over scaled pseudotime along each of the trajectories from (E). **(G)** Loess curves fitted to the expression of selected genes and surface proteins over scaled pseudotime along each of the trajectories from (E). **(H)** Dot plot of selected genes identified by differential analysis (p < 0.05, |log_2_(FC)| ≥ 0.9) between the early and late phases of each trajectory. The size of each ball represents the proportion of cells with non-zero expression, colour represents fold-change at early or late phases. Early <=120 DPI, late > 120 DPI. **(I)** Dot plot of enrichment scores from GSEA using differentially expressed genes between early and late stages within each trajectory independently. Normalised enrichment scores (NES) were calculated independently within each trajectory. Positive NES indicates enrichment of gene set early in the trajectory, and negative NES indicates late enrichment. Results were considered significant (Sig.) if p < 0.05.

**Fig. S5.**
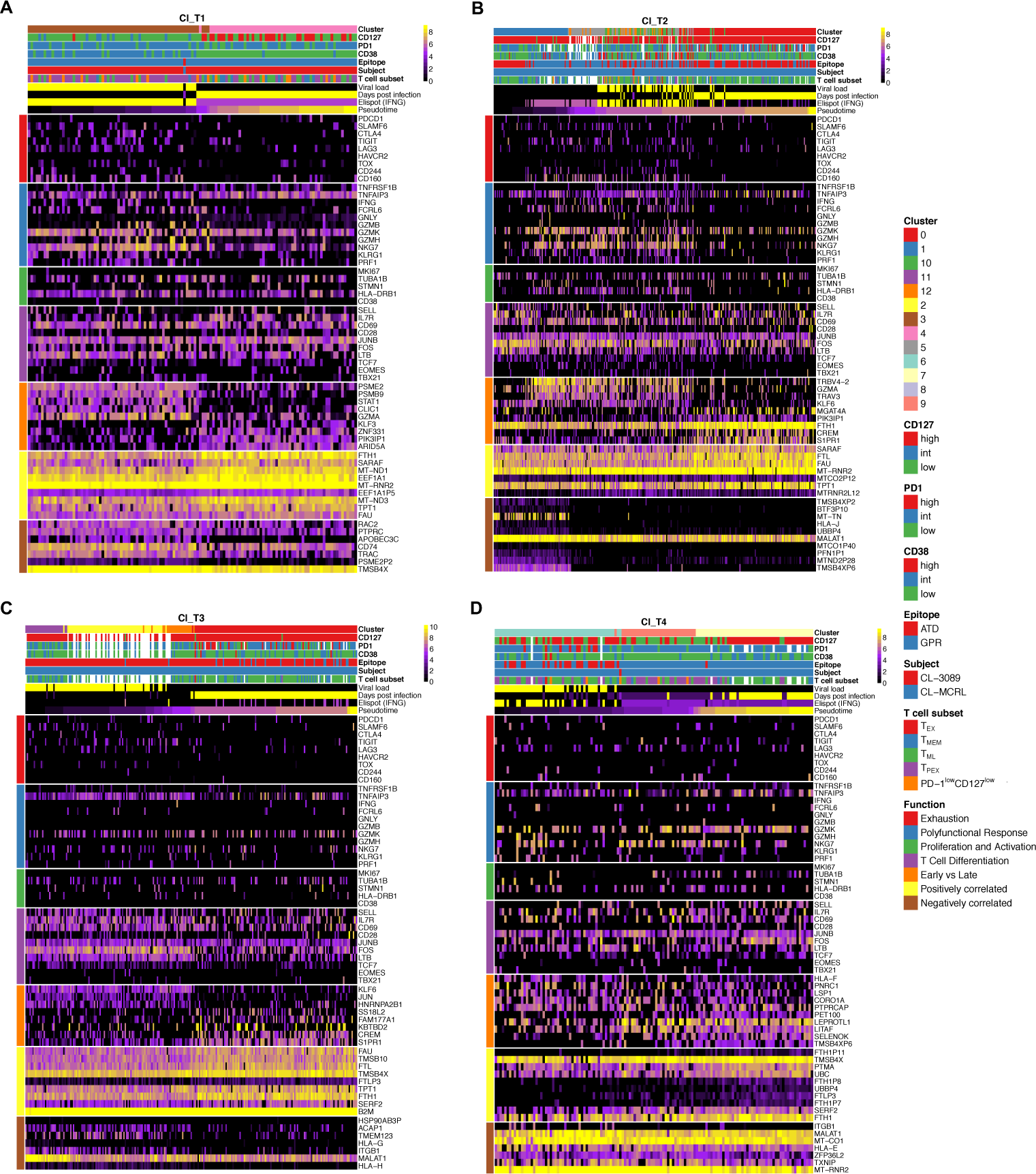
Phenotypic and molecular evolution of HCV-specific T cells associated with trajectories identified from clearers. **(A-D)** Heatmaps of representative genes from cells forming trajectories Cl_T1 **(A)**, Cl_T2 **(B)**, Cl_T3 **(C)** and Cl_T4 **(D).** Cells are ordered by pseudotime revealing positive correlation with the Days post infection. For each heatmap metadata are reported on T-cell phenotype (index sorting), epitope specificity, subject of origin, viral load and IFN-γ ELISpot measured at the sample of origin. Selected genes from differential expression analysis between early and late phase of infection, as well as genes correlated with pseudotime were utilised for these plots. Genes were grouped based on their functions. T_EX_/exhausted: PD-1^high^CD127^low^, T_PEX_/progenitor-exhausted: PD-1^int^CD127^low^, T_ML_/memory-like: PD-1^high^CD127^high^, T_MEM_/memory: PD-1^low^CD127^high^.

**Fig. S6.**
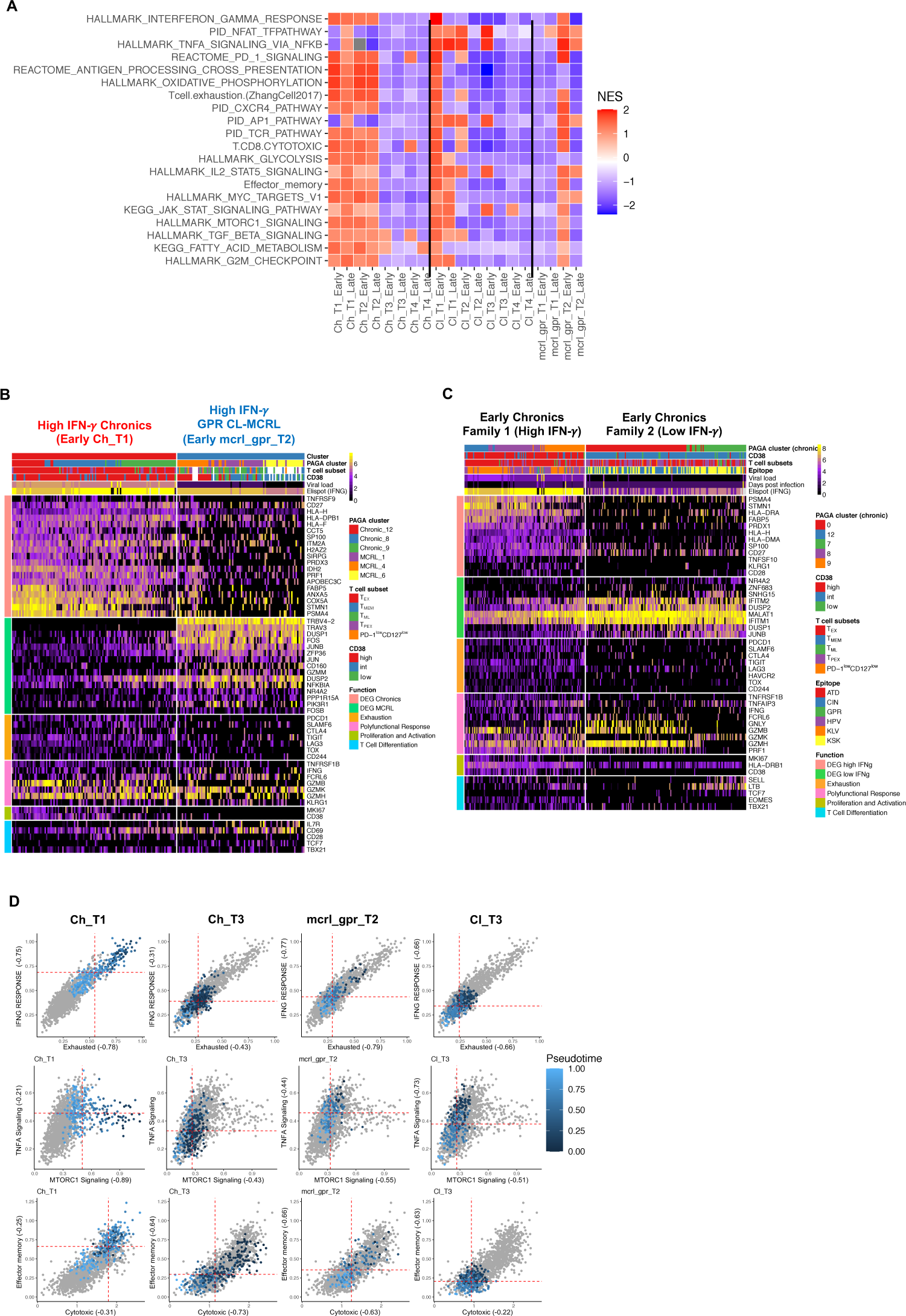
Trajectory analysis reveals the molecular and phenotypic features of HCV-specific T cells in acute phase of infection that explain magnitude of the response. **(A)** GSEA heatmap of enriched pathways identified between the early (≤120 DPI) and late (>120 DPI) phases of each trajectory. High NES indicates enrichment of pathway. Significant (in at least one trajectory phase with p < 0.05) pathways shown are selected from Hallmark, PID, KEG and Reactome gene sets. NES: Normalised enrichment score. **(B)** Heatmap of gene expression for differentially expressed genes between early cells from the family 1 trajectories in chronic progressors, and early cells in the mcrl_gpr_T2 trajectory associated with early cytotoxic function and IFN-*γ* production. Significant (p < 0.05) genes with the largest fold change between the two clusters are shown, in addition to genes known to be functional markers. Cells are annotated their T cell subset and CD38 expression. **(C)** Heatmap of gene expression for differentially expressed genes with the largest fold changes between early phase (≤120 DPI) cells from chronic family 1 (high IFN-γ) and family 2 (low IFN-γ) trajectories combined with genes from a curated list. Cells are annotated according to T cell subset, epitope and CD38 expression. **(D)** Scatter plots depicting gene signatures score (mean value) across trajectories. Each dot represents a cell, and cells belonging to the same trajectory are colour coded based on pseudotime value. Correlation (Pearson’s) between gene set and pseudotime are shown on axis if significant (p < 0.05). Red, dashed-line represents mean gene score of the cells in the trajectory.

## Supplementary tables

**Table S1.** Clinical and laboratory characteristics of the subjects (N=17) included in this study.

| Subjects | Age at infection /Sex | Symptomatic | HLA type |  |  | HCV genotype | No. of days post infection <sup>a</sup> | HCV Ab | HCV RNA | Viral Load (IU/ml) | Number of epitopes detected |
| --- | --- | --- | --- | --- | --- | --- | --- | --- | --- | --- | --- |
|  |  |  | A | B | C |  |  |  |  |  |  |
| CH-240 | 21/M | Yes | 0201 | 1501 | 0304 | 3a | -61 | - | - | 0 |  |
|  |  |  | 0201 | 5701 | 0602 |  | 44 | - | + | 54,887 | 3 |
|  |  |  |  |  |  |  | 57 | + | + | 85,473 |  |
|  |  |  |  |  |  |  | 71 | + | + | 64,063 | 3 |
|  |  |  |  |  |  |  | 85 | + | + | 6,051 | 3 |
|  |  |  |  |  |  |  | 99 | + | + | 497 | 3 |
|  |  |  |  |  |  |  | 113 | + | + | 4,862 | 3 |
|  |  |  |  |  |  |  | 140 | + | + | 1,034 | 3 |
|  |  |  |  |  |  |  | 220 | + | + | 44,449 | 3 |
|  |  |  |  |  |  |  | 310 | + | + | 29,552 | 3 |
|  |  |  |  |  |  |  | 538 | + | + | 62,174 | 3 |
| CH-023 | 22/M | Yes | 0201 | 4402 | 0501 | 1a | -165 | - | - | 0 |  |
|  |  |  | 0201 | 5701 | 0602 |  | 36 | - | + | 19,234,348 | 8 |
|  |  |  |  |  |  |  | 44 | - | + | 17,907,338 | 8 |
|  |  |  |  |  |  |  | 60 | + | + | 8,121,396 | 8 |
|  |  |  |  |  |  |  | 74 | + | + | 397,185 | 8 |
|  |  |  |  |  |  |  | 85 | + | + | 3,218 |  |
|  |  |  |  |  |  |  | 101 | + | + | 398 | 8 |
|  |  |  |  |  |  |  | 135 | + | + | 2,843,176 | 8 |
|  |  |  |  |  |  |  | 197 | + | + | 5,896,155 | 8 |
|  |  |  |  |  |  |  | 365 | + | + | 516,945 | 8 |
|  |  |  |  |  |  | 3a | 635 | + | + | 14,738 | 8 |
|  |  |  |  |  |  |  | 781 | + | + | 344,388 |  |
| CH-HOKD | 26/F | Yes | 0101 | 0801 | 0602 | 1b | -135 | - | - | 0 |  |
|  |  |  | 3001 | 1302 | 0701 |  | 30 | - | + | 733,849 |  |
|  |  |  |  |  |  |  | 65 | NA | NA |  |  |
|  |  |  |  |  |  |  | 72 | + | + | 175,219 |  |
|  |  |  |  |  |  |  | 79 | + | + | 44,452 | 5 |
|  |  |  |  |  |  |  | 93 | + | + | 407,392 |  |
|  |  |  |  |  |  |  | 107 | + | + | 24,969 | 5 |
|  |  |  |  |  |  |  | 121 | + | + | 77,723 |  |
|  |  |  |  |  |  |  | 149 | + | + | 254,245 | 5 |
|  |  |  |  |  |  |  | 170 | + | NA |  |  |
|  |  |  |  |  |  |  | **233** | + | + | 350,658 | 5 |
|  |  |  |  |  |  | 1a | 446 | + | + | 956,488 |  |
|  |  |  |  |  |  |  | 618 | + | + | 102,439 |  |
|  |  |  |  |  |  |  | 985 | + | + | 14,587 |  |
| CH-256 | 30/M | No | 0301 | 0702 | 0401 | 1a |  | - | - | 0 |  |
|  |  |  | 2402 | 3501 | 0702 |  | **44** | - | + | 34,149,824 |  |
|  |  |  |  |  |  |  | **58** | + | + | 19,188,762 | 6 |
|  |  |  |  |  |  |  | **79** | + | + | 812,622 | 6 |
|  |  |  |  |  |  |  | **96** | + | + | 50,774 |  |
|  |  |  |  |  |  |  | 112 | + | + | 50 | 6 |
|  |  |  |  |  |  |  | 128 | + | + | 693 |  |
|  |  |  |  |  |  |  | 162 | + | + | 135 | 6 |
|  |  |  |  |  |  |  | **286** | + | + | 14,853 | 6 |
|  |  |  |  |  |  |  | 300 | + | + | 732 |  |
|  |  |  |  |  |  |  | 569 | + | + | 16,421 |  |
|  |  |  |  |  |  |  | 944 | + | + | 85,835 |  |
| CH-THDS | 25/M | No | 0201 | 1402 | 0102 | 1a | -152 | - | - | 0 |  |
|  |  |  | 3201 | 2705 | 0802 |  | **16** | - | + | 235,662 |  |
|  |  |  |  |  |  |  | 30 | - | + | 549,251 |  |
|  |  |  |  |  |  |  | 44 | - | + | 176,550 |  |
|  |  |  |  |  |  |  | 58 | + | + | 2,005 |  |
|  |  |  |  |  |  |  | 72 | + | + | 108,737 | 3 |
|  |  |  |  |  |  |  | 85 | + | + | 62,043 |  |
|  |  |  |  |  |  |  | **109** | + | + | 503,981 |  |
|  |  |  |  |  |  |  | **198** | + | + | 681,389 |  |
|  |  |  |  |  |  |  | 395 | + | + | 223,849 |  |
|  |  |  |  |  |  |  | 530 | + | + | 183,426 |  |
| CH-684MX | 27/M | Yes | 0201 | 2702 | 0202 | 1a | -166 | - | - | 0 |  |
|  |  |  | 0301 | 4001 | 0304 |  | **2** | - | + | 140,200 |  |
|  |  |  |  |  |  |  | 16 | - | + | 98,867 |  |
|  |  |  |  |  |  |  | 30 | - | + | 78,764 |  |
|  |  |  |  |  |  |  | 44 | - | + | 33,763 |  |
|  |  |  |  |  |  |  | **58** | + | + | 19,932 |  |
|  |  |  |  |  |  |  | 71 | + | + | 20,374 | 5 |
|  |  |  |  |  |  |  | 95 | + | + | 28,181 |  |
|  |  |  |  |  |  |  | **184** | + | + | 221,964 |  |
|  |  |  |  |  |  |  | 380 | + | + | 75,688 |  |
|  |  |  |  |  |  |  | 515 | + | + | 143,746 |  |
| CL-360 | 28/M | Yes | 3201 | 1402 | 0501 | 3a | -110 | - | - | 0 |  |
|  |  |  | 6802 | 4402 | 0802 |  | **30** | - | + | 5,648,631 |  |
|  |  |  |  |  |  |  | 44 | - | + | 4,617,483 | 3 |
|  |  |  |  |  |  |  | 58 | + | + | 14,170 |  |
|  |  |  |  |  |  |  | 71 | + | + | 15,938 | 3 |
|  |  |  |  |  |  |  | 83 | + | + | 1,060 |  |
|  |  |  |  |  |  |  | 97 | + | + | <15 | 3 |
|  |  |  |  |  |  |  | 132 | + | + | 57 |  |
|  |  |  |  |  |  |  | 223 | + | - | 0 |  |
|  |  |  |  |  |  |  | 422 | + | - | 0 | 3 |
| CL-277 | 24/M | No | 0201<br>1101 | 4402<br>4402 | 0501<br>0501 | 3a | -578 | - | - | 0 |  |
|  |  |  |  |  |  |  | 39 | - | + | 5,482,503 |  |
|  |  |  |  |  |  |  | 63 | + | + | 12,442,419 |  |
|  |  |  |  |  |  |  | 74 | + | + | 10,374,554 |  |
|  |  |  |  |  |  |  | 95 | + | + | 3,473,088 |  |
|  |  |  |  |  |  |  | 102 | + | + | 10,506 |  |
|  |  |  |  |  |  |  | 116 | + | + | 70,120 | 6 |
|  |  |  |  |  |  |  | 144 | + | + | 833 |  |
|  |  |  |  |  |  |  | 245 | + | - | 0 |  |
|  |  |  |  |  |  |  | 437 | + | - | 0 |  |
| CL-MCRL | 25/F | No | 0101<br>2902 | 0702<br>4403 | 0702<br>1601 | 1a | -81 | - | - | 0 |  |
|  |  |  |  |  |  |  | 80 | + | + | 1,846 |  |
|  |  |  |  |  |  |  | 87 | + | + | 58 | 2 |
|  |  |  |  |  |  |  | 115 | + | - | 0 | 2 |
|  |  |  |  |  |  |  | 171 | + | - | 0 |  |
|  |  |  |  |  |  |  | 256 | + | - | 0 | 2 |
|  |  |  |  |  |  |  | 487 | + | - | 0 |  |
|  |  |  |  |  |  |  | 648 | + | - | 0 | 2 |
|  |  |  |  |  |  |  | 732 | + | - | 0 |  |
| CL-087 | 32/F | NA | 2402<br>3004 | 1402<br>1506 | 0403<br>0802 | 1b | -46 | - | - | 0 |  |
|  |  |  |  |  |  |  | 31 | + | + | 13,118,082 |  |
|  |  |  |  |  |  |  | 42 | + | + | 29,257,428 |  |
|  |  |  |  |  |  |  | 61 | + | + | 6,455,009 |  |
|  |  |  |  |  |  |  | 70 | + | + | 1,940,469 |  |
|  |  |  |  |  |  |  | 116 | + | + | 25 | 3 |
|  |  |  |  |  |  |  | 133 | + | - | 0 |  |
| CL-364 | 29/M | Yes | 0101<br>0301 | 0702<br>5701 | 0602<br>0702 | 1a | -337 | - | - | 0 |  |
|  |  |  |  |  |  |  | 337 | + | + | 1,932 | 2 |
|  |  |  |  |  |  |  | 352 | + | - | 0 | 2 |
|  |  |  |  |  |  |  | 629 | + | - | 0 | 2 |
| CL-231 | 22/M | NA | 0101 | 0702 | 0602 | 3a | **57** | + | + | 2,242,163 | 1 |
|  |  |  | 0101 | 5701 | 0702 |  | 199 | + | - | 0 | 1 |
| CL-089 | 26/M | No | 0101 | 0702 | 0501 | 1b | -180 | - | - | 0 |  |
|  |  |  | 3001 | 4402 | 0702 |  | **180** | + | + | 70,737 | 1 |
|  |  |  |  |  |  |  | 356 | + | - | 0 | 1 |
|  |  |  |  |  |  |  | 534 | + | - | 0 | 1 |
|  |  |  |  |  |  |  | 752 | + | - | 0 | 1 |
| CI-101 | 36/M | NA | 0101 | 0801 | 0501 | 1a/3a | -536 | - | - | 0 |  |
|  |  |  | 0201 | 4402 | 0701 |  | 178 | + | + | 11441811 | 2 |
|  |  |  |  |  |  |  | 215 | + | + | 7646402 | 2 |
|  |  |  |  |  |  |  | 333 | + | + | 0 | 2 |
|  |  |  |  |  |  |  | 385 | + | + | 0 | 2 |
| CH-221 | 21/M | NA | 1101 | 0702 | 0102 | 3a | 46.5 | + | + | 315892 | 1 |
|  |  |  | 3001 | 4601 | 0702 |  | 523.5 | + | + | 200219 |  |
|  |  |  |  |  |  |  | 787.5 | + | + | 294540 |  |
| CH-132 | 20/F | NA | 0101 | 0801 | 0303 | 3a | -262.5 | - | - | 0 | 1 |
|  |  |  | 3101 | 5501 | 0701 |  | 262.5 | + | + | 26678 |  |
|  |  |  |  |  |  |  | 472.5 | + | + | 65365 |  |
|  |  |  |  |  |  |  | 691.5 | + | + | 118153 |  |
|  |  |  |  |  |  |  | 842.5 | + | + | 164179 |  |
|  |  |  |  |  |  |  | 1564.5 | + | + | 73491 |  |
| CH-059 | 30/M | NA | 0101 | 0702 | 0701 | 1a | -80.5 | - | - | 0 | 1 |
|  |  |  | 0201 | 0801 | 0702 |  | 13.5 | - | + | 3676682 |  |
|  |  |  |  |  |  |  | 27.5 | - | + | 17409803 |  |
|  |  |  |  |  |  |  | 41.5 | NA | + | 26187596 |  |
|  |  |  |  |  |  |  | 60.5 | + | + | 7308585 |  |
|  |  |  |  |  |  |  | 76.5 | + | + | 488992 |  |
|  |  |  |  |  |  |  | 88.5 | + | + | 5906 |  |
|  |  |  |  |  |  | 2 | 102.5 | + | + | 588467 |  |
|  |  |  |  |  |  |  | 195.5 | + | + | 1096633 |  |
|  |  |  |  |  |  |  | 423.5 | + | + | 321276 |  |

<sup>a</sup> The number of days post infection was estimated from the time to seroconversion.
Bold denotes time points with viral samples sequenced for detection of autologous virus using Next Generation Sequencing.

**Table S2.** Viral sequencing and epitopes detected and tested in ELISPOT assays.

| *Subject* | **Outcome of infection** | **Geno-type** | **Viral genome sequence (Sanger)** | **Deep Sequencing (NGS)** | **Number of epitopes tested in ELISPOT\*** |
| --- | --- | --- | --- | --- | --- |
| *CH-240* | Chronic | 3a | + | + | 70 |
| *CH-023* | Chronic | 1a | + | + | 45 |
| *CH-HOKD* | Chronic | 1b | + | + | 82 |
| *CH-256* | Chronic | 1a | + | + | 98 |
| *CH-THDS* | Chronic | 1a | + | + | 99 |
| *CH-684MX* | Chronic | 1a | + | + | 100 |
| *CL-360* | Clearer | 3a | + | + | 93 |
| *CL-277* | Clearer | 3a | + | + | 69 |
| *CL-MCRL* | Clearer | 1a | + | - | 3 |
| *CL-087* | Clearer | 1b | + | + | 71 |
| *CL-364* | Clearer | 1a | + | + | 8 |
| *CL-231* | Clearer | 3a | + | + | 41 |
| *CL-089* | Clearer | 1b | + | - | 5 |
| *CL-101* | Clearer | 1a,3a | + | - | 9 |
| *Ch-221* | Clearer | 3a | - | - | 1* |
| *Ch-059* | Chronic | 1a | + | + | 1* |
| *Ch-132* | Chronic | 3a | - | - | 0 |

\*Tested with negative results.

**Table S3:**
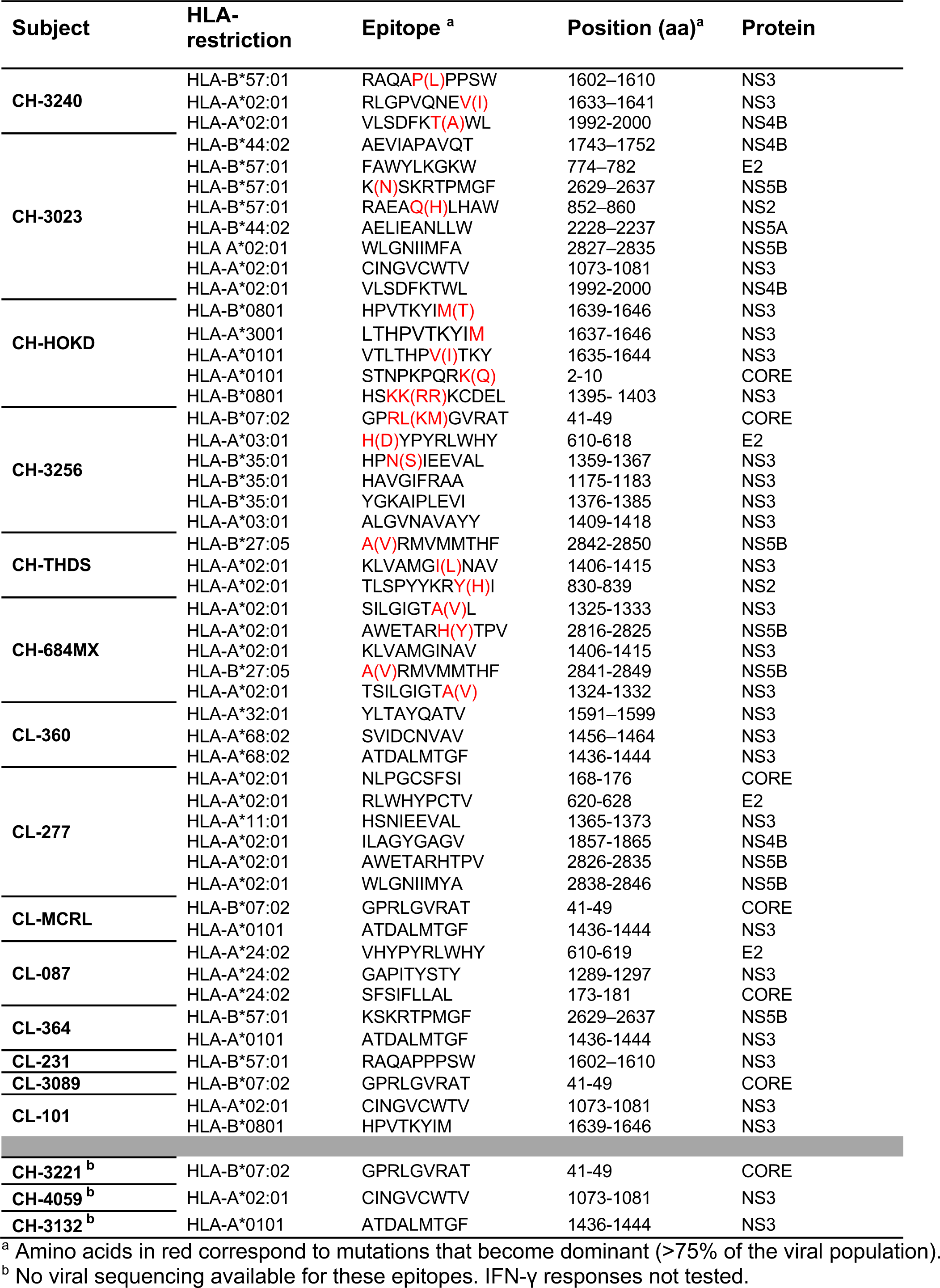
HLA-I restricted T-cell epitopes identified with positive IFN-γ ELISpot responses for each subject.

**Table S4:** HLA class I dextramers used for immunophenotyping of HCV-specific CD8^+^ T cells.

| HCV protein | Position (aa) | Epitope sequence | WT/escape | T cells identified | HLA-I | Subjects |
| --- | --- | --- | --- | --- | --- | --- |
| NS3 | NS3 1602–1610 | RAQAPPPSW | WT | Y | B*57:01 | CH-3240; CL-231 |
| NS3 | 1602–1610 | RAQALPPSW | ESCAPE | N | B*57:01 | CH-3023 |
| NS3 | 1633–1641 | RLGPVQNEV | WT | Y | A*02:01 | CH-3240 |
| NS3 | 1633–1641 | RLGPVQNEI | ESCAPE | N | A*02:01 | CH-3240 |
| NS4B | 1992-2000 | VLSDFKTWL | WT | Y | A*02:01 | CH-3023; CH-3240 |
| NS4B | 1992-2000 | VLSDFKAWL | ESCAPE | N | A*02:01 | CH-3023; CH-3240 |
| NS5B | 2629–2637 | KSKRTPMGF | WT | Y | B*57:01 | CH-3023; CL-364 |
| NS5B | 2629–2637 | NSKRTPMGF | ESCAPE | N | B*57:01 | CH-3023 |
| NS2 | 852–860 | RAEAQLHAW | WT | N | B*57:01 | CH-3240 |
| NS3 | 1073-1081 | CINGVCWTV | WT | Y | A*02:01 | CH-3023; CH-4059; CL-101 |
| NS3 | 1639-1646 | HPVTKYIM | WT | Y | B*0801 | CL-101;CH-HOKD |
| NS3 | 1395- 1403 | HSKKKCDEL | WT | N | B*0801 | CH-HOKD |
| E2 | 610-618 | HYPYRLWHY | WT | Y | A*03:01 | CH-3256 |
| CORE | 41-49 | GPRLGVRAT | WT | Y | B*07:02 | CH-HOKD;CH-3256;CH-3221;CL-3089 |
| NS3 | 1359-1367 | HPNIEEVAL | WT | Y | B*35:01 | CH-3256 |
| NS3 | 1175-1183 | HAVGIFRAA | WT | N | B*35:01 | CH-3256 |
| NS3 | 1406-1415 | KLVMGINAV | WT | N | A*02:01 | CH-684MX |
| NS3 | 1436-1444 | ATDALMTGF | WT | Y | A*0101 | CL-364;CL-3132; CL-MCRL |
| NS3 | 1436-1444 | ATDALMTGY | ESCAPE | N | A*0101 | CL-364 |
| NS5B | 2827–2835 | WLGNIIMFA | WT | N | A*02:01 | CH-3023 |

**Table S5:** Estimates of the rate of escape for viral epitopes.

| Subject ID | Epitope (escape variant)* | Epitope (WT) | IFN-γ ELISPOT<sup>&</sup> | Rate of escape (Std Error) | P-value |
| --- | --- | --- | --- | --- | --- |
| CH-HOKD | HPVTKYIT | HPVTKYIM | 2425 | 0.387 (0.074) | 0.002 |
|  | LTHPVTKYIT | LTHPVTKYIM | 1385 | 0.387 (0.074) | 0.002 |
|  | HSRRKCDEL | HSKKKCDEL | 55 | 0.245 (0.027) | 0.0001 |
| CH-THDS | KLVMGLNAV | KLVMGINAV | 1992.5 | 0.330 (0.037) | 0.0009 |
|  | TLSPYYKRHI | TLSPYYKRYI | 67.5 | 0.085 (0.014) | 0.0036 |
|  | VRMVMTHF | ARMVMMTHF | 655 | 0.160 (0.074) | 0.0700 |
| THGS0684MX | TSILGIGTV | TSILGIGTA | 230 | 0.350 (0.372) | 0.4000 |
|  | SILGIGTVL | SILGIGTAL | 997.5 | 0.350 (0.213) | 0.4000 |
|  | AWETARYTPV | AWETARHTPV | 50 | 0.080 (0.008) | 0.0087 |
| CH-3256 | DYPYRLWHY | HYPYRLWHY | 750 | 0.560 (0.039) | 0.0007 |
|  | GPKMGVRAT | GPRLGVRAT | 195 | 0.047 (0.003) | 0.0050 |
|  | HPSIEEVAL | HPNIEEVAL | 150 | 0.049 (0.002) | 0.0000 |
| CH-3023 | RAEAHLHAW | RAEAQLHAW | 261 | 0.080 (0.005) | 0.0001 |
|  | NSKRTPMGF | KSKRTPMGF | 854 | 0.166 (0.035) | 0.0090 |
| CH-3240 | RAQALPPSW | RAQAPPSW | 220 | 0.102 (0.016) | 0.0080 |
|  | RLGPVQNEI | RLGPVQNEV | 380 | 0.003 (0.002) | 0.0007 |

\* Dominant escape variant (i.e., frequency of occurrence >70%)
<sup>&</sup> Maximum value of SFU per million PBMC against wild type epitope measured within the first 120DPI.

**Table S6:** TCR detection (includes VDJPuzzle TCR reconstruction from scRNA-seq data, and sanger sequencing)

| Subject | Epitope | Number of cells with TCR$\alpha\beta$ (%) | Unique CDR3$\alpha\beta$ | Expanded CDR3$\alpha\beta$ clones* | Number of cells with CDR3α | Number of unique CDR3α | Number of expanded CDR3α clones | Number of cells with CDR3β | Number of unique CDR3β | Number of expanded CDR3β clones |
| --- | --- | --- | --- | --- | --- | --- | --- | --- | --- | --- |
| CL-MCRL | GPR | 174 (71%) | 2 | 2 | 181 | 2 | 2 | 228 | 3 | 2 |
| CL-MCRL | ATD | 262 (80%) | 129 | 46 | 275 | 111 | 45 | 304 | 129 | 59 |
| CL-3089 | GPR | 98 (75%) | 44 | 16 | 100 | 41 | 17 | 125 | 42 | 23 |
| CH-THDS | KLV | 116 (97%) | 22 | 10 | 116 | 21 | 10 | 118 | 21 | 10 |
| CH-HOKD | HPV | 99 (92%) | 63 | 16 | 100 | 52 | 19 | 107 | 65 | 19 |
| CH-4059 | CIN | 83 (70%) | 59 | 11 | 88 | 59 | 12 | 105 | 75 | 15 |
| CH-3256 | HYP | 37 (55%) | 4 | 1 | 38 | 4 | 1 | 66 | 2 | 1 |
| CH-3256 | GPR | 14 (78%) | 5 | 3 | 14 | 5 | 3 | 18 | 4 | 2 |
| CH-3240 | RLG | 143 (99%) | 22 | 7 | 144 | 19 | 7 | 143 | 10 | 7 |
| CH-3240 | RAQ | 123 (98%) | 64 | 13 | 126 | 55 | 13 | 123 | 13 | 8 |
| CH-3221 | GPR | 63 (70%) | 10 | 4 | 65 | 9 | 4 | 87 | 12 | 5 |
| CH-3132 | ATD | 26 (62%) | 20 | 4 | 30 | 23 | 4 | 36 | 28 | 4 |
| CH-3023 | CIN | 130 (65%) | 117 | 6 | 137 | 118 | 11 | 186 | 162 | 11 |
| CH-3023 | KSK | 195 (71%) | 100 | 38 | 198 | 96 | 40 | 267 | 86 | 42 |

\* Number of CDR3 $\alpha\beta$ clones that are observed more than once

The following tables are uploaded separately

**Table S7:** Metadata file of single cells, including full length TCR sequences.

**Table S8:** Differentially expressed genes between groups of cells or trajectories.

**Table S9:** Customised gene signatures utilised for the GSEA.

**Table S10:** ELISpot data for each epitope specific CD8^+^ T cells.

